# Hierarchical importance of factors impacting trilobite cephalic disparity

**DOI:** 10.1101/2025.07.08.663686

**Authors:** Harriet B. Drage, Stephen Pates

## Abstract

Trilobite cephala show high disparity owing to their supporting of key functional ecological structures, such as sensory and feeding structures, and those enabling important behaviours like enrollment and exoskeleton moulting. Previous studies analysed associations between cephalic morphometry, trends through time and with diversity, and these behaviours. However, no studies have yet quantitatively compared the importance of these different varied factors on cephalic morphometry. Here, we explore the association between trilobite cephalic morphometry and taxonomic variables, moulting behaviour, and facial suture type in Principal Components Analysis morphospace, and assess whether these variables can predict cephalic morphometry using Linear Discriminant Analyses. We employed Linear Mixed-Effects Models (LMMs) to quantify the hierarchical importance of these variables for determining cephalic disparity. The dataset used comprises cephalon outlines for 762 species of trilobites from across the globe and through the entire Palaeozoic. We found that variation in cephalic morphometry is not clearly associated with moulting behaviour, nor is moulting behaviour linked to facial suture type. Trilobite moulting behaviour is evidently too phenotypically plastic, and/or preservationally impacted, to produce clear signals across broad datasets. Results demonstrate the utility of LMMs for palaeontological datasets; we found that family-level assignment overall well explains cephalic morphometry, more so than order-level assignment which shows significant overlap in morphospace. Geological Period, facial suture type, and moulting behaviour had little impact on cephalic morphometry. We thereby demonstrate that trilobite cephalic outline morphometry is generally in accordance with trilobite family-level systematics and highlight the use of LMMs for interrogating broad-scale fossil morphological data.

## 2 Introduction

### Trilobite morphological disparity

Trilobites, the most abundant and speciose animal group in the Palaeozoic, were incredibly important parts of early ecosystems, occupying oceans globally and filling a wide diversity of ecological niches. Across their long evolutionary history, trilobites showed high and varying levels of disparity in different aspects of their morphology (e.g., Foote, 1993; Hopkins, 2014, 2022; Cole and Hopkins, 2021; Suárez and Esteve, 2021; Bault et al., 2023; Holmes, 2023; see summary in Drage and Pates, 2024). In particular, the cephalon shows high disparity owing to its supporting of key features for ecological function, including sensory organs, feeding structures, coaptive structures for enrollment behaviour, and the facial sutures and other moulting structures (Fortey and Owens, 1999; Drage et al., 2018a; Suárez and Esteve, 2021). This manifests in the huge variety of cephalic phenotypes seen in trilobites and can be seen in the diversity of overall shapes of these cephala. Analysing the trajectories of cephalic shape change provides crucial information on how the disparity of trilobite form, and their occupation of ecological niches, changed through the Palaeozoic across periods of dramatic global change such as during the early Cambrian and Ordovician (Foote, 1991, 1997; Webster, 2007; Hopkins, 2014; Esteve et al., 2021b; Holmes, 2023; Drage and Pates, 2024). Exploring cephalic shape change also provides a method to study trilobite behavioural evolution on a broad scale; for example, links between cephalic shape and enrollment strategy (Suárez and Esteve, 2021), and association with exoskeleton moulting behaviour (Drage, 2019, 2024; Drage et al., 2023).

While there have been numerous analytical studies of trilobite morphometry (see summary in Drage and Pates, 2024), which test the associations between morphometry of the cephalon or pygidium with various behaviours and other abiotic and biotic variables (e.g., Hopkins, 2014), none have quantitatively compared the relative importance of the different potential drivers of cephalic morphometry. Many interacting aspects shape our view of the trilobite cephalic phenotype, including ecological and behavioural factors like feeding mode, moulting mode, locomotory method, and enrolment type, as well as taxonomic assignment because we partially base our systematic scheme on cephalic morphology, and geological age of occurrence because trilobite species are limited to the morphospace range occupied at the time in which they diverged. We thereby apply Principal Components Analysis, Linear Discriminant Analysis, and Linear Mixed-Effects Modelling to explore the hierarchical importance of drivers of trilobite cephalic morphometry. To that end, we will answer:

1. Is taxonomic assignment, particularly to family level, the best predictor of trilobite occupation of cephalon outline morphospace, or are other variables related to evolutionary history important drivers of cephalic shape?
2. Do trilobite species that moulted in different ways have significantly different cephalon outline shapes, and can cephalon shape be used to predict moulting behaviour?
3. Is there a link between the facial suture type and moulting behaviour a trilobite species showed?

### Trilobite moulting and morphology

Moulting is a complex process that all arthropods, including trilobites, must repeat throughout their lives to grow and develop. Drage et al. (2018a) and Drage (2019) showed that trilobite moulting behaviour is both interspecifically and intraspecifically variable, which we might expect to be due to variation in morphology. The presence or absence of the facial sutures, cephalic planes of weakness that functioned to facilitate moulting (Du et al., 2024) though may have also been structural (Esteve et al., 2021b), led to dramatic differences in the moulting behaviours employed, however, less direct morphological differences presumably also impacted the trilobite moulting process. For example, changes in size and shape might have driven intraspecific variability in moulting, which broad morphological differences like facial suture presence cannot explain. However, studies have found little link between moulting behaviour and traditional morphometry (the relative sizes and shapes of different tagma) (Drage et al., 2023; Drage, 2024), except for the outright presence or absence of functional facial sutures during different ontogenetic stages of the same species (Drage et al., 2018b). Drage et al. (2023) found that, from a large study of *Estaingia bilobata*, the species has high intraspecific variability in moulting behaviour and the configurations preserved, but that this had only a minor link to differences in body proportions.

Traditional morphometry seemingly thereby fails to explain the interspecific and intraspecific variability in trilobite moulting behaviour. However, the cephalic exoskeleton is arguably the most crucial part of the trilobite exoskeleton in terms of moulting. This is because its morphology enables all major modes of moulting (Henningsmoen, 1975; Drage, 2019), which involve either sutures on the cephalon (variously the facial sutures, rostral suture, hypostomal suture, marginal suture, ventral suture), or opening of the cephalothoracic joint articulation. Studies have yet to interrogate whether the shape of the cephalon itself has any bearing on the type of moulting behaviour observed. Further, trilobites with functional facial sutures display a range of suture types, which were historically (though generally no longer) used to group them taxonomically (Stubblefield, 1936). No studies have so-far quantitatively assessed whether facial suture type has any bearing on moulting behaviour, other than their overall presence/absence (Drage, 2019).

## 3 Materials and methods

### Dataset

We used the dataset presented in Drage and Pates (2024), with a single specimen representing each of 419 species, and combined with the published dataset of Suárez and Esteve (2021) (343 species with agnostids removed); totalling 762 species for the entire dataset. These species each have outline semilandmark coordinates for two curves (of 64 coordinates, total 128 coordinates), representing the anterior margin and posterior margin of the cephalon (see Drage and Pates, 2024). Where represented in the raw data of Drage and Pates (2024) by multiple specimens, a single representative specimen was chosen that lay between the range of coordinates for the species (i.e., the closest real specimen to that of the median). Each of the species have taxonomic assignments to family level based on those in Adrain (2011) and geological Period occurrences using the original descriptive literature and Jell and Adrain (2002). One order assignment for the orders and families was ‘Uncertain’, pertaining to families previously assigned to the ‘waste-basket’ order Pytchopariida (see Adrain, 2011). We also recorded the facial suture type of all species through checking morphology in the original descriptive literature (see raw data in Supplementary I); this includes opisthoparian, gonatoparian, proparian, marginal (around the anterior margin of the cephalon), ventral (ventral cephalic sutures opening, as in Olenellidae; Henningsmoen, 1975; Palmer and Repina, 1993), and anterior median (at the median of the ventral side of the cephalon, as in asaphids; Henningsmoen, 1975; Whittington, 2003; Zhu et al., 2010).

We also used a subset of this dataset focused around exoskeleton moulting mode, with data on moulting behaviour for available species taken from the datasets published by Drage (2019, 2024). Each of the 216 species for which moulting data were available were summarised as showing one of the following moulting behaviour categories: Sutural Gape, with opened facial sutures and disarticulated librigenae; Ventral Gape, with opened ventral sutures (rostral and potentially hypostomal) and a disarticulated rostral plate where present; Salter’s, with an opened cephalothoracic joint and disarticulated cephalon; Henningsmoen’s (after Henningsmoen’s configuration; Drage et al., 2018a), where both the facial sutures and cephalothoracic joint are opened together in the same specimen; and Both, where some specimens of a species show the Sutural Gape category and others the Salter’s category (but not in the same specimen as for Henningsmoen’s). Other ecological factors beyond moulting mode, such as feeding and life mode, may also impact cephalic morphometry. However, data from the Paleobiology Database (Uhen et al., 2023), the online database containing global trilobite occurrence data, is not at the required resolution to address this question (see Supplementary IV).

### Analyses

Geometric morphometric analyses and multivariate statistical analyses were performed on the two datasets described above (total dataset and moulting subset) (Fig. 1). All analyses were carried out in R (version #4.4.0 Puppy Cup; R Core Team, 2024) using Rstudio (Posit Team, 2025) (code provided in Supplementary II). A total dataset morphospace and a moulting subset morphospace were produced using elliptical Fourier analysis (EFA) with 13 harmonics that represent 99.9% of the variation retained, as in (Drage and Pates, 2024). Principal Components Analyses (PCA) using the package momocs (Bonhomme et al., 2014) were used to visualise relevant patterns of shape variation and analyse differences in morphospace occupation between taxonomic family groups, facial suture groups, and moulting mode groups.

**Figure 1:**
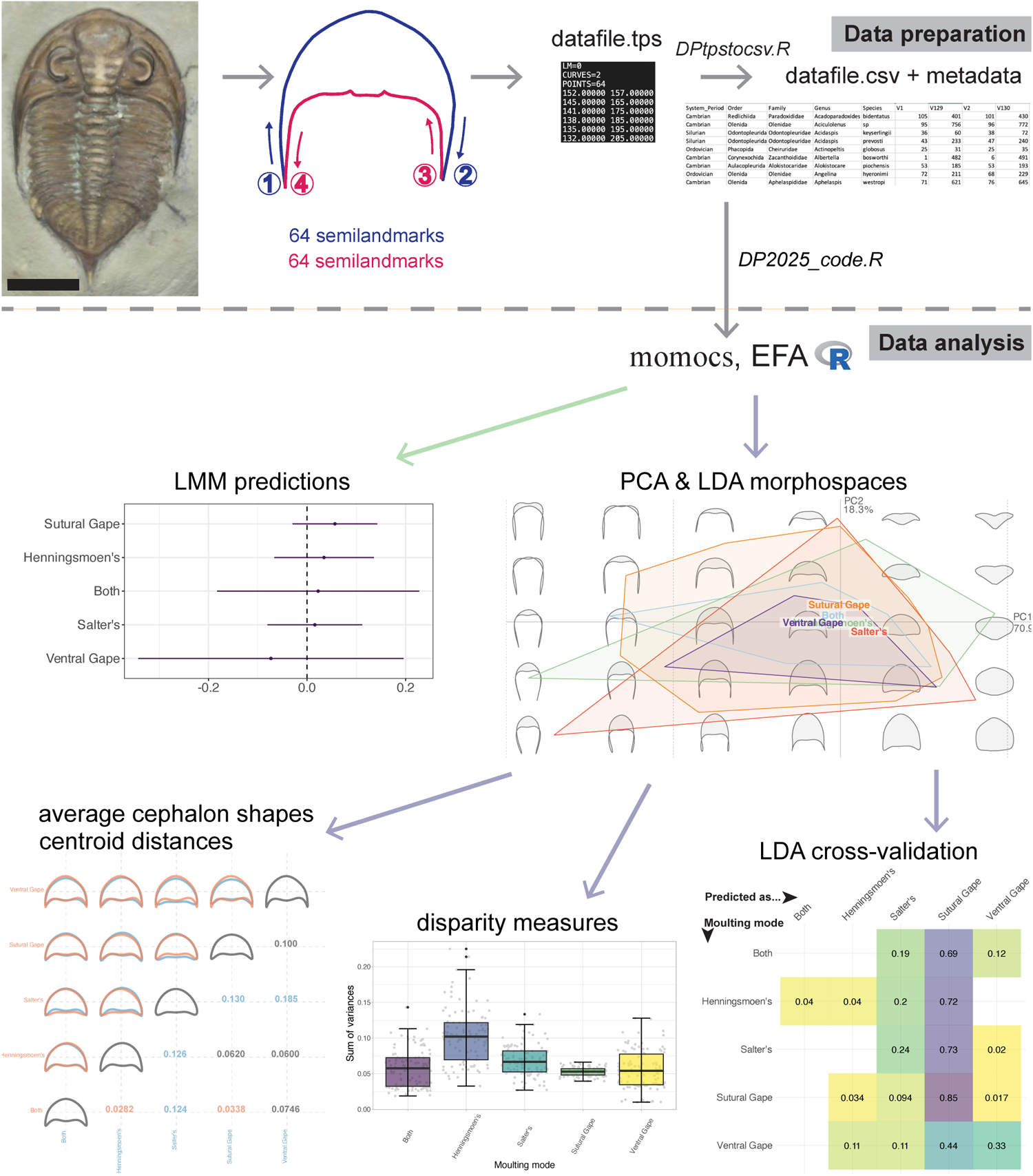
Analytical pathway. Data preparation section from Drage and Pates (2024). Code file provided as Supplementary II.

Groupings in PCA morphospace were further interrogated by calculating the convex hull areas (except for families as there were too many groups) and comparing the group centroid distances and average cephalon outline shapes. Multivariate Analysis of Variance analyses (MANOVA) were performed to test for differences in the distributions of principal components (PC) scores between groupings. For most downstream analyses using family groupings, families with n = <5 were removed, leaving 42 families across 12 orders. Only PCs 1 and 2 were plotted and used for subsequent analyses; the morphospace scree plots show that these are responsible for 64.6% and 24.5% of the variation respectively for the total morphospace, and 70.9% and 18.3% respectively for the moulting data subset. Summed PCs from 3 onwards therefore represent ∼10% of the total variation in morphospace occupation.

Linear Discriminant Analyses (LDA) were performed using the momocs package (Bonhomme et al., 2014) to produce cross-validation tables. These LDA cross-validation tables inform on the probability that a new data point within each group would be placed within each other group (i.e., what is the probability of its category being predicted correctly or incorrectly). This therefore provides an estimation of the predictiveness of cephalic outline for each grouping in the dataset. The LDAs also allow plotting of linear discriminants in morphospace, which aim to separate the groupings as much as is possible.

Disparity measures (sums of variances, SoV, and sums of ranges, SoR) were calculated for all groupings using the dispRity package (Guillerme, 2018). SoV describes the density of points in morphospace; a low SoV indicates specimens are more densely clustered in morphospace. SoR is a measure of total morphospace occupation and is more sensitive to sample size and outliers; a lower SoR means a smaller area of morphospace occupied (Hopkins, 2022). The disparity measures were compared using pairwise t-tests (alpha level of 0.05 with Bonferroni correction = 0.05/n pairings) to determine whether the disparities significantly differ between the groups (see full results in Supplementary III).

A linear regression analysis was performed (using the packages tidyverse and broom; Wickham et al., 2019) to determine to what extent moulting mode and facial suture group covary in the dataset. We would expect some covariance in cephalon outline shape between these variables (i.e., some comparable morphological adaptation, and therefore position in morphospace) because moulting mode is at least partially determined by facial suture type. For example, the Sutural Gape mode of moulting requires dorsal facial sutures (opisthoparian, gonatoparian, or proparian) and the Ventral Gape mode requires ventral sutures. However, not all moulting modes are clearly related to facial suture type.

Finally, Linear Mixed-Effects Model (LMM) analyses were performed using the package lme4 (Bates et al., 2015) to analyse the impact of all categorical variables on cephalon outline morphometry (PC1 and PC2 coordinates). Altering the choice of these variables as fixed effects or random effects allowed for determination of their hierarchical importance on placement in morphospace. Fixed effects in LMMs denote variables of interest, that is, the independent variables we want to test the impact of on the dependent variable (here, the PC1 or PC2 coordinate in morphospace). Random effects denote the other variables that we want to control the influence of. Thus, the result for a given LMM fixed effect estimates how much the PC coordinate of focus would change given a one unit change in the fixed effect variable. The best-fitting model with each categorical variable separately designated as the fixed effect was considered that with the lowest (most negative) Aikake Criterion (AIC), with the log likelihood values (highest being better fitting) also checked but are more vulnerable to sampling. The effect sizes of the best-fitting LMMs were then assessed by their conditional R^2^ (cR^2^; the total proportion of variability in the dependent variable [PC coordinate] explainable by the whole model) and marginal R^2^ (mR^2^; the proportion of variability in the dependent variable explainable only by the model fixed effect). The magnitudes of the effect sizes were compared for each fixed effect variable, and the overall effect size of the fixed effect assessed for significance. Speekenbrink (2023) and Wiley (2020) provide particularly useful explanations of all components of LMM analyses. The package lme4 does not provide p-value estimates (see Bates et al., 2015, and Luke, 2017, for reasoning), though this can be obtained for the fixed effect overall, along with more detailed model information, from the packages multilevelTools and JWileymisc (Wiley, 2020, 2025). The full LMM results are available in Supplementary III.

The LMMs were constructed with the following format:

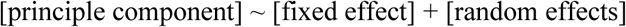

For the random effects, these were variously tested as (1|random effect) being just the random intercepts, and (fixed effect|random effect) being the random intercepts and random slopes for each level of the random effect. These random effects also had their interactions included as (random effect) + (random effect) being just the main effects, (random effect) * (random effect) being the main effects and interactions of the variables, or certain random effects were excluded from the model entirely. Different order polynomials were also tested for the fixed effects as poly(random effect, [polynomial order]). The total list of variables tested (individually as fixed effects, and incorporated as random effects, as above) is order, family, genus, species, geological Period, facial suture type, and moulting mode. LMMs were not tested with genus and species as fixed effects because they can be assumed to have the biggest impact on position in cephalon outline shape morphospace because species descriptions are, at least partially, based on cephalon morphology and so the model reasoning would be circular. Both PC1 and PC2 were individually tested as the dependent variable, that is, the variable changing in response to the fixed effect. For most model tests polynomials decreased the fit, and testing the random intercepts (without the random slopes; (1|random effect)) improved model fit, was less computationally intensive (the random slopes occasionally being too intensive), and made the models less complex (which is preferred by the lme4 package; Bates et al., 2015). All models were run on a 2020 MacBook Pro 2.3 GHz Quad-Core Intel Core i7 with 16 GB RAM.

## 4 Results

### Family assignment

Drage and Pates (2024) demonstrated significant association between cephalon outline shape and order taxonomic assignment but noted that some orders have high disparities that may be obscuring finer-scale morphometric or evolutionary trends. Interrogating these cephalon outline data at the family-level does show clear differences in the locations and areas of PCA morphospace occupation, both within and between the orders (Fig. 2). In general, each family clusters in a single, reasonably restrained area in morphospace. Some families show notably higher morphospace areas than others, particularly the Asaphidae, Saukiidae, Harpetidae, Lichidae, Odontopleuridae, Olenidae, Acastidae, Cheiruridae, Proetidae, Raphiophoridae and Trinucleidae. Other families appear to have singular outliers, potentially artificially extending their area of morphospace occupied and suggesting they otherwise have lower disparities – this includes the Styginidae, Harpetidae, Acastidae, and to a lesser extent the Olenidae, Olenellidae and Conocoryphidae (Fig. 2). However, several diverse and well-sampled groups show comparably low areas of morphospace occupation, notably the Phacopidae and Calymenidae – the similarly diverse and well-sampled Proetidae occupy a larger area than either of these families. Several families cluster around the centre of morphospace, including the Asaphidae, Dolichometopidae, Dorypygidae, Asaphiscidae, Acastidae, Cheiruridae, Dalmanitidae, Phacopidae, Pterygometopidae, Proetidae, and Ellipsocephalidae. Others are positioned at the extremes of cephalon outline morphospace occupation, such as the Cyclopygidae and Nileidae at positives of PC1, Harpetidae at the negative of PC2, and Lichidae somewhat to the positive of PC2 (Fig. 2). A significant MANOVA result (Table 1) supports the family groups differing in their occupation of PCA morphospace, although the low F value suggests these differences are not extensive.

**Figure 2:**
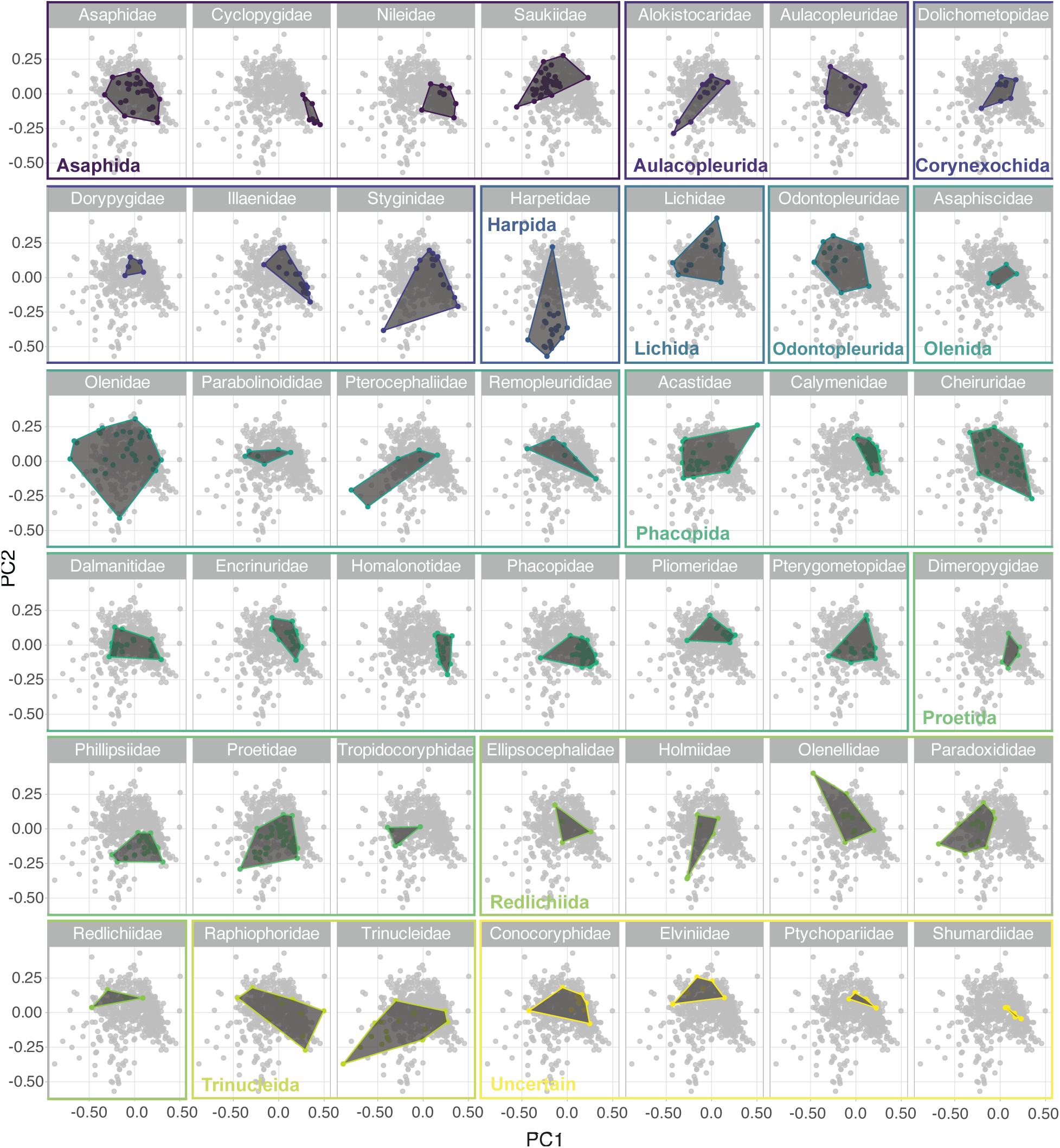
Principal Components Analysis of the dataset subdivided by family groups. Each facet has the family displayed named at the top, with plotted species belonging to that family coloured on a background of the total dataset (in grey dots), connected by the convex hull. The colours and coloured, labelled boxes display to which order each family belongs.

**Table 1:** Sample size, PCA convex hull area, and MANOVA results. Family group sample sizes in Supplementary I.

| **Moulting mode** | **Convex hull area** | **Sample size (# species)** | **Facial suture type** | **Convex hull area** | **Sample size** |
| --- | --- | --- | --- | --- | --- |
| Both | 0.150 | 16 | Gonatoparian | 0.240 | 62 |
| Henningsmoen's | 0.355 | 25 | Marginal | 0.485 | 55 |
| Salter's | 0.423 | 49 | Opisthoparian | 0.681 | 430 |
| Sutural Gape | 0.446 | 117 | Proparian | 0.356 | 183 |
| Ventral Gape | 0.133 | 9 | Ventral | 0.385 | 24 |
|  |  |  | Anterior median | 0.0148 | 8 |
| **MANOVA group** | **F value** | **p value** |  |  |  |
| Moulting mode | 2.87 | $8.19 \times 10^{-08}$ | Retained to 9 PCs | | |
| Facial suture type | 11.8 | $<2.20 \times 10^{-16}$ | Retained to 10 PCs | | |
| Family | 3.98 | $<2.20 \times 10^{-16}$ | Retained to 10 PCs | | |

The family SoRs correspond roughly with morphospace area occupied, so the families with high SoRs (Fig. 3) are those that occupy the largest area, including Trinucleidae (SoR = 1.607), Raphiophoridae (1.401), Olenidae (1.722), Acastidae (1.213), Harpetidae (1.224), and Proetidae (1.030). However, several high SoRs are presumably artificially inflated by their apparent outliers, such as for Styginidae (1.399) and Olenellidae (1.169) (full results in Supplementary III). The well-sampled families with low SoRs also generally correspond with those with low morphospace areas, such as Homalonotidae (SoR = 0.485), Cyclopygidae (0.407), Calymenidae (0.566) and Encrinuridae (0.635). These families with low SoRs generally also have notably low SoVs (Fig. 3), including the Calymenidae, Homalonotidae, Phacopidae, Encrinuridae, Pliomeridae, Phillipsiidae, and Saukiidae (all >10 species sample size and <0.025 SoV) (Fig. 3, Supplementary I and III). This indicates extensive clustering within morphospace for these families, that is, they are predominantly restricted to one area and have overall low disparity. With particularly high SoVs (still >10 species) are the Trinucleidae (SoV = 0.116), Raphiophoridae (0.103), Olenidae (0.093), Acastidae (0.057), and Styginidae (0.066) (Fig. 3, Supplementary III). This suggests less clustering in these families, with more extensive spread over the area of morphospace occupied, and high disparities for these groups given their additionally high SoRs. The majority of family pairwise t-tests were significant with Bonferroni correction for both SoV and SoR, including for those families discussed above; the full results are given in Supplementary III.

**Figure 3:**
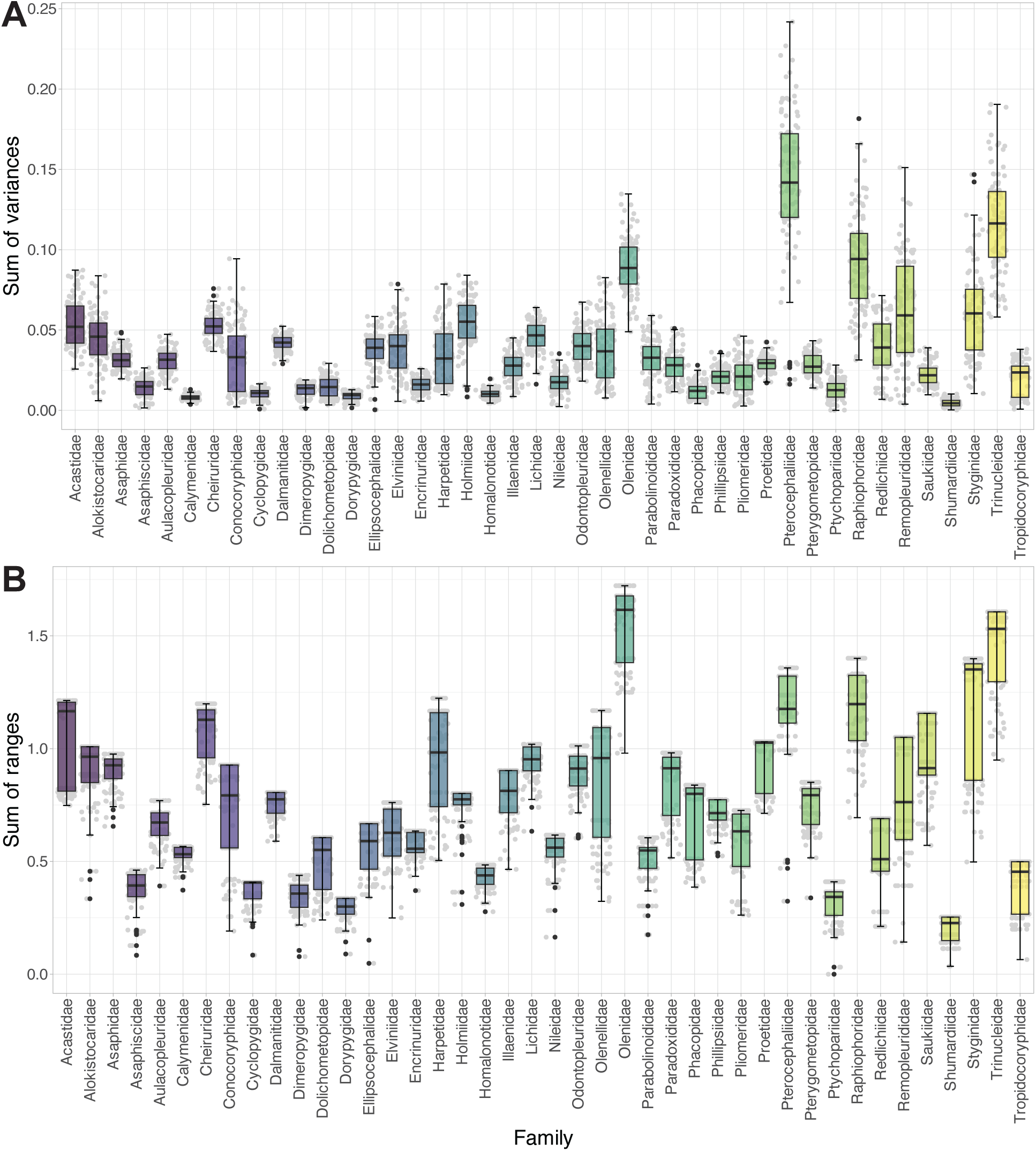
A, sum of variances box and whisker plots for family groups of the Principal Components Analysis; B, sum of ranges box and whisker plots of the same. Black lines show the median values, hinges correspond to the first and third quartiles, and the error bars represent 1.5× the interquartile range. Black dots are those species that fall outside the error bars (i.e., potential outliers), and the grey dots display all underlying data points (horizontal placement of points is meaningless).

Pairwise PCA centroid distance comparisons for all families are available in Supplementary III, with there being too many pairings to fully display here. Some family centroids are extremely close (<0.01 distance), notably the Cheiruridae and Dolichometopidae, and Illaenidae with Calmoniidae and Dokimocephalidae, though there are many other families that are also very close in morphospace at a distance slightly exceeding 0.01 (Supplementary III). Many other family pairings are quite distant (>0.3 distance), and a range of families have high distances from many other families. The Harpetidae are notably distant from all other families (distance from closest family >0.2), with distances >0.5 from many (including many phacopid families; Calmoniidae, Calymenidae, Conocoryphiidae, and Encrinuridae). Several other families have distances >0.3 from most other families, including Trinucleidae, Paradoxididae, Alsataspididae, Bathycheilidae, and Cyclopygidae. Supplementary Figure 1 displays the average cephalon outline shapes for all families, demonstrating extensive differences between family cephalic outline (particularly for those already noted as distant to most other families, e.g., Harpetidae); these differences are much greater than the differences between the representative orders (see Drage and Pates, 2024, fig. 5), explaining their particularly high pairwise centroid distances.

Few family assignments would be correctly predicted by LDA using cephalon outline morphometry alone (Supplementary Figure 2). Interestingly, the LDA cross-validation table gives few strong probabilities across all family groups; any predictions, correct or incorrect, are difficult to make based on cephalon outline shape alone. This indicates that the morphospace is not being dominated by a few families (unlike for the order Phacopida; see Drage and Pates, 2024), but also that there is sufficient overlap (likely due to the high number of groupings tested) to prevent families being correctly predicted. The most likely families to be predicted for a new species are the Calymenidae, Cheiruridae, Olenidae, Phacopidae, Saukiidae, and somewhat Proetidae. However, these families do not necessarily have the highest SoRs or areas of morphospace occupation (except Olenidae), or all cluster at the morphospace origin (Fig. 2). It might be, therefore, that this result is more related to the sample size of these families as they are amongst the highest (Supplementary I), though not exclusively. Most families do not have a high probability of being predicted correctly; only the Cheiruridae, Elviniidae, Olenidae, Phacopidae, and Pharostomatidae have a higher or equal probability of being correctly predicted than assigned to any other singular family, but these probabilities are still low (usually p << 0.5). This seems unrelated to the morphospace area these families occupy as they do not have the highest SoRs, but is presumably related to morphospace positioning and convex hull overlap as these families are generally centrally positioned (Fig. 2). Some families that appear to be at the extremes of occupied morphospace (Fig. 2), such as the Harpetidae, would still not be correctly predicted using LDA.

### Moulting behaviour

The PCA results plotted in morphospace for the moulting mode groupings demonstrate extensive overlapping between the convex hulls to the point that they are almost entirely nested (Fig. 4A). The centroids of each group are also similar, with all centroids clustering around the centre of morphospace. The Sutural Gape convex hull occupies a particularly broad area of morphospace (area = 0.45; Table 1) that ranges across the total region of morphospace occupied by the dataset (Fig. 4). The Salter’s group convex hull is also high in area (0.42) though clustered further right at more positive values of PC1 and spread across the extent of occupied PC2 (Fig. 4C). The other moulting mode groups are also somewhat clustered towards the centre of morphospace (Fig. 4C), though occupying smaller morphospace areas (Table 1). The Ventral Gape convex hull occupies the smallest area (0.13) but has the smallest sample size. The few points at the most negative extreme of PC1 for at least the Salter’s and Henningsmoen’s groups may be outliers artificially extending the morphospace area occupation (Fig. 4C). A significant MANOVA result (Table 1) suggests that the moulting mode groups differ in their occupation of morphospace on PCs 1 and 2, though the F value is comparably low, suggesting these differences are minor.

**Figure 4:**
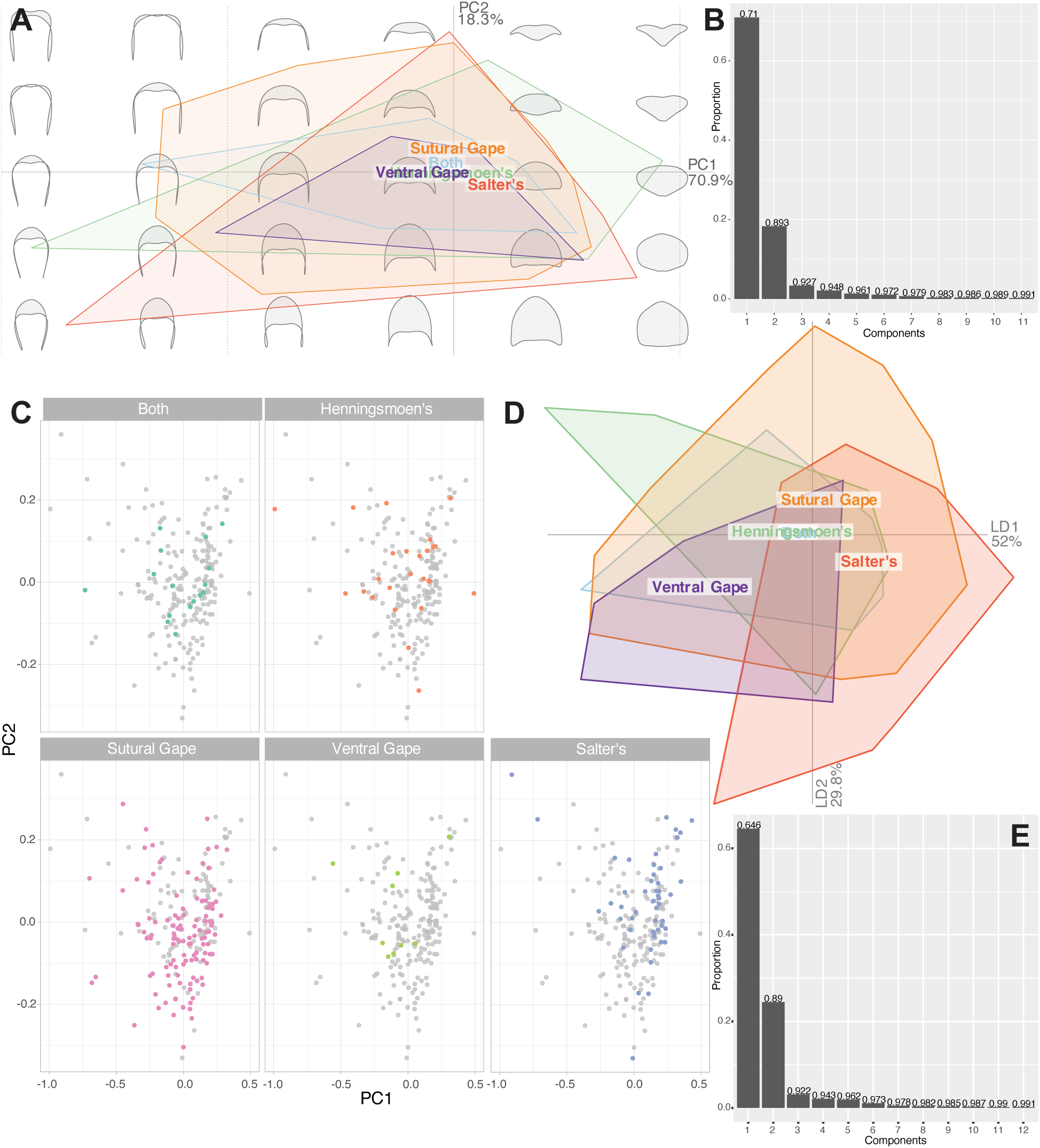
A, Principal Components Analysis (PCA) for moulting sub-dataset, with all moulting mode convex hulls displayed on the same morphospace; B, scree plot for PCA of moulting sub-dataset (demonstrating that PCs 1 and 2 represent ∼89% of the dataset variance); C, same data as for A, but with each moulting mode highlighted separately in colour on a separate morphospace (grey points represent the total dataset); D, Linear Discriminant Analysis of moulting sub-dataset; E, scree plot for the entire dataset. Some group labels in A and D overlap because they represent the relevant centroid positions.

The pairwise centroid distance results help to quantify the variation in PCA morphospace occupation, and demonstrate similarities between the cephalon outline shapes of some groups that are difficult to interpret from the PCA plots (Fig. 5A). These distances suggest similar cephalon outline morphometries in the Both—Sutural Gape—Henningsmoen’s groups (pairwise distances <0.07), with the Henningsmoen’s group also being close to the Ventral Gape group (0.6). The other two moulting groups are comparably distant from all other groups. This weak morphogroup and two singular moulting type groups is clear in the similar average cephalon shapes (Fig. 5A). The Salter’s group has an average cephalon that is axially longer with shorter genal spines, and the Ventral Gape group is also axially long but with pronounced genal spines, while the Both, Sutural Gape, and Henningsmoen’s average shapes are very similar to each other (Fig. 5A).

**Figure 5:**
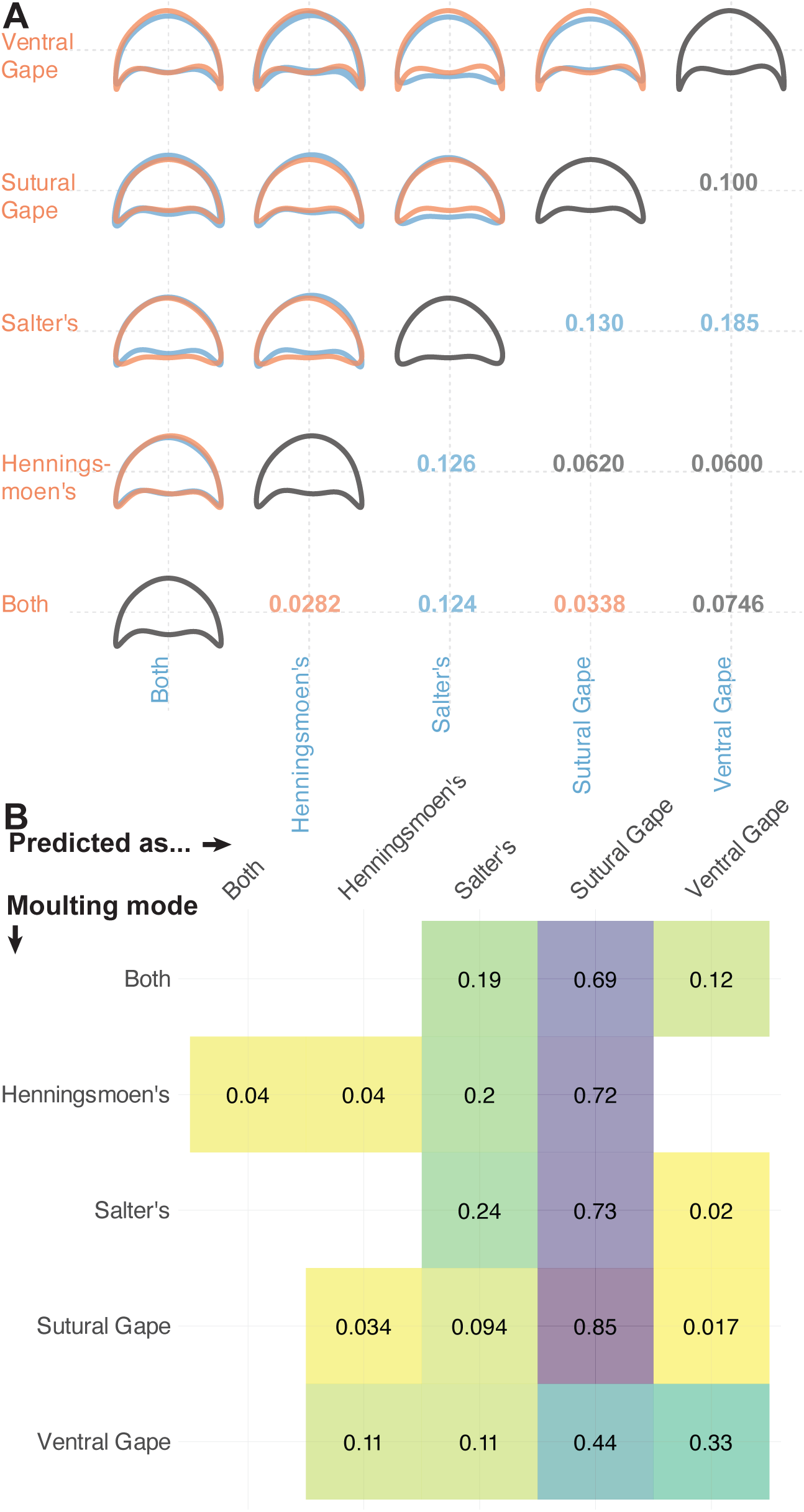
A, Pairwise comparison of average cephalic shape for the moulting mode groups, and pairwise centroid distances in Principal Components Analysis morphospace between the groups. Notably close centroid pairs (<0.05) are in orange text, and notably distant centroid pairs (>0.1) are in blue text. B, Linear Discriminant Analysis cross-validation table. The table gives the probability that a species with a given true moulting mode (y-axis) will be predicted as each of the moulting modes along the x-axis. Empty cells equate to a negligible (close to zero) probability.

The LDA plot (Fig. 4D) demonstrates some ability of cephalon outline shape to distinguish the moulting mode groups in morphospace, resulting in less overlap of groups than in the PCA morphospace (Fig. 4A). Of note are the movement of the Salter’s group hull and centroid away from the centre of morphospace to more positive values of LD axis 1 and of the Ventral Gape to more negative values of both LD axes. The Sutural Gape centroid is also positioned more positive on LD axis 2, while Both and Henningsmoen’s retain overlapping centroids close to the morphospace centre.

The LDA cross-validation table (Fig. 5B) suggests that cephalon outline morphometry has little capability for predicting the moulting behaviour of a trilobite species. A new species added to the dataset is most likely to be placed in the Sutural Gape moulting mode, regardless of the species’ true moulting behaviour, as a result of the group’s broad and central occupation of trilobite morphospace (Figs 4, 5B). For example, Salter’s group species show a 0.24 probability of species having their moulting behaviour correctly identified, but a 0.73 probability of being confused with the Sutural Gape mode. A true Ventral Gape species is the least likely to be predicted as a Sutural Gape mode specimen (p = 0.44), though this is still the most likely outcome for the Ventral Gape group with a correct prediction chance of only 0.33.

The SoVs (Fig. 6A) are similar across the moulting mode groups (Fig. 6A), with the groups showing extensive overlap in their error bars. Only for the Henningsmoen’s group is the SoV particularly higher (0.102, all others <0.072), suggesting a slightly greater spread of specimens in morphospace for this group. The SoRs for the Henningsmoen’s, Salter’s and Sutural Gape modes are all reasonably high (1.95, 2.03 and 1.614 respectively; Fig. 6B), indicating comparably broader areas of morphospace occupation, and corresponding with their high convex hull areas. For the Henningsmoen’s group, the high SoV and SoR suggest an overall relatively high disparity, while the Sutural Gape and Salter’s groups occupy a broad area but show only moderate disparity as their lower SoVs indicate most species are more tightly clustered within the occupied morphospace. The ubiquitously significant pairwise t-test results after Bonferroni correction (Supplementary III) support the Henningsmoen’s group SoV being significantly higher than the other groups. The only other significant pairing is between the SoVs for Salter’s and Sutural Gape moulting modes. For the SoR, all but the Henningsmoen’s–Sutural Gape and Both–Ventral Gape pairings are significant (Supplementary III).

**Figure 6:**
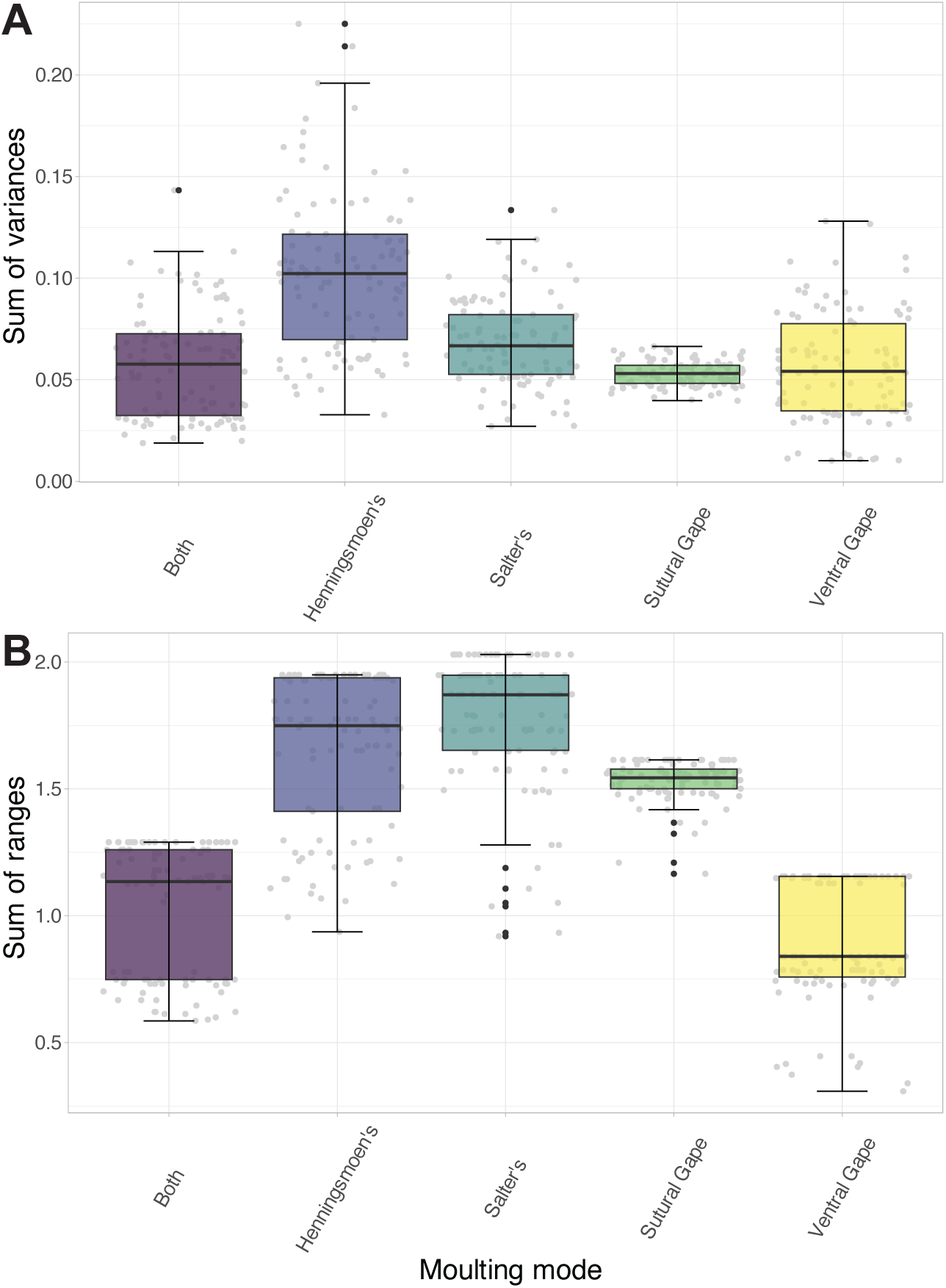
A, sum of variances box and whisker plots for moulting mode groups of the Principal Components Analysis; B, sum of ranges box and whisker plots of the same. Black lines show the median values, hinges correspond to the first and third quartiles, and the error bars represent 1.5× the interquartile range. Black dots are those species that fall outside the error bars (i.e., potential outliers), and the grey dots display all underlying data points (horizontal placement of points is meaningless).

### Facial suture type

The groups representing different cephalic suture types are significantly different in their occupation of morphospace plotted using PCA (Fig. 7), with the corresponding significant MANOVA F value being high (Table 1); this suggests more divergent cephalic shapes for the facial suture groups than the moulting mode groups. There remains extensive overlap of the cephalic suture groups around the centre of the PCA morphospace (Fig. 7A), though the extents of the convex hulls and some centroids differ in position. The anterior median suture group is at the extreme positive end of PC1, and slightly negative on PC2, with a small convex hull area (0.0148) reflected by their small sample size (Table 1). The marginal suture group has a centroid negative on PC1 and 2 and with a convex hull extending to the extremes of PC2; all other group centroids have positive PC2 positions. The marginal suture group has the second highest convex hull area (0.485), with only opisthoparian being larger (0.681). The other suture types all have comparable convex hull areas (0.240–0.385), with centroids slightly positive on PC2 but close to the centre, and slightly positive (proparian, gonatoparian) or negative (opisthoparian, ventral) on PC1 (Fig. 7A). Some clustering differences are evident between the groups (Fig. 7B). For example, despite the medium-sized convex hull for the gonatoparian suture group, almost all specimens of the group are clustered at positive PC1 values, separated only along the PC2 axis; this suggests the extreme points may be outliers. Similarly, while the opisthoparian convex hull has the largest area, the part at negative PC1 and PC2 values is sparsely populated compared to the large region around the centre of morphospace (Fig. 7B). The ventral group also shows some specimens isolated at the extremes, though the marginal group specimens show no real clustering nor obvious outliers. The proparian group appears to show one major cluster at positive PC1 differentiated along PC2, on a background of many species spread across broader values of PC1 and 2.

**Figure 7:**
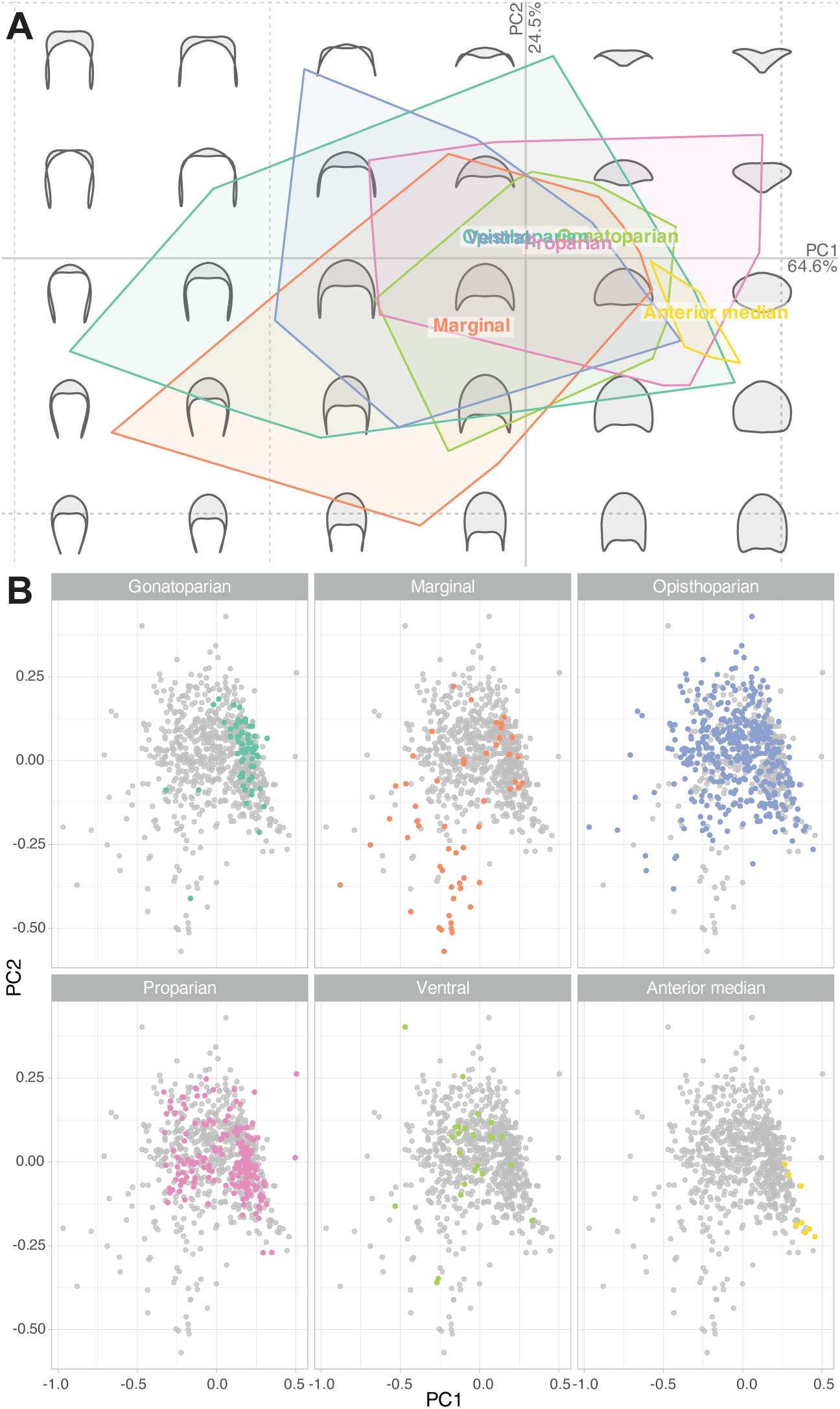
A, Principal Components Analysis (PCA) for total dataset grouped by facial suture type, with all convex hulls displayed on the same morphospace; B, same data as for A, but with each facial suture group highlighted separately in colour on a separate morphospace (grey points represent the total dataset). Some group labels in A overlap because they represent the relevant centroid positions.

The LDA plot (Fig. 8A), unlike for the moulting mode groups (Fig. 4D), does not show any clear additional group differentiation in morphospace occupation compared to the PCA morphospace (Fig. 7A). The convex hulls show even more overlap at the centre of the morphospace (even the anterior median group), with relatively isolated specimens at the extremes producing the furthest extents of the large convex hull areas. The group centroids also show more overlap at the centre of the LDA morphospace.

**Figure 8:**
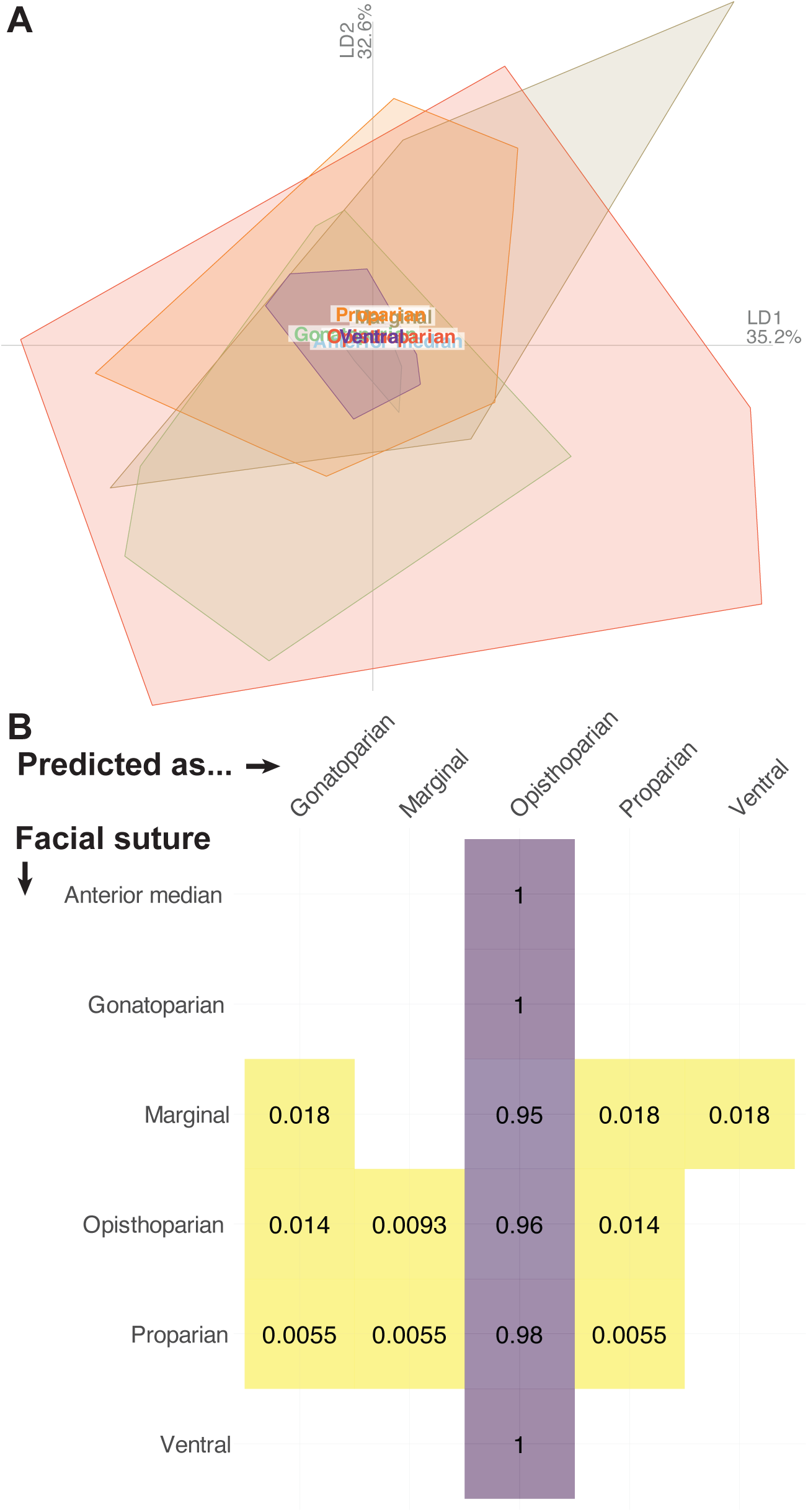
A, Linear Discriminant Analysis (LDA) of facial suture groups in total dataset; B, LDA crossvalidation table. The table gives the probability that a species with a given true facial suture type (y-axis) will be predicted as each of the types along the x-axis. Empty cells equate to a negligible (close to zero) probability. Group labels in A overlap because they represent the relevant centroid positions.

The pairwise centroid distances and average cephalon outline shapes for the facial suture types reveal some cephalic outline similarities between different groups (Fig. 9). The proparian centroid is close to the gonatoparian, opisthoparian, and, to a lesser extent, the ventral suture groups, though the average proparian cephalon differs in its lack of genal spines. The ventral suture and opisthoparian groups show almost identical average cephala (and very close centroids) that are axially short with medium genal spines (Fig. 9), though the ventral suture group has a reasonably low sample size (Table 1). The anterior median suture group has an almost circular average cephalon outline morphometry with no genal protrusions that is very different, and distant, to all other groups (Fig. 9), though again the group has a low sample size. Similarly, the marginal suture group has a divergent average cephalon and centroid position distant from the other groups, due to its deep axial shape with longer and broader genal spines (Fig. 9).

**Figure 9:**
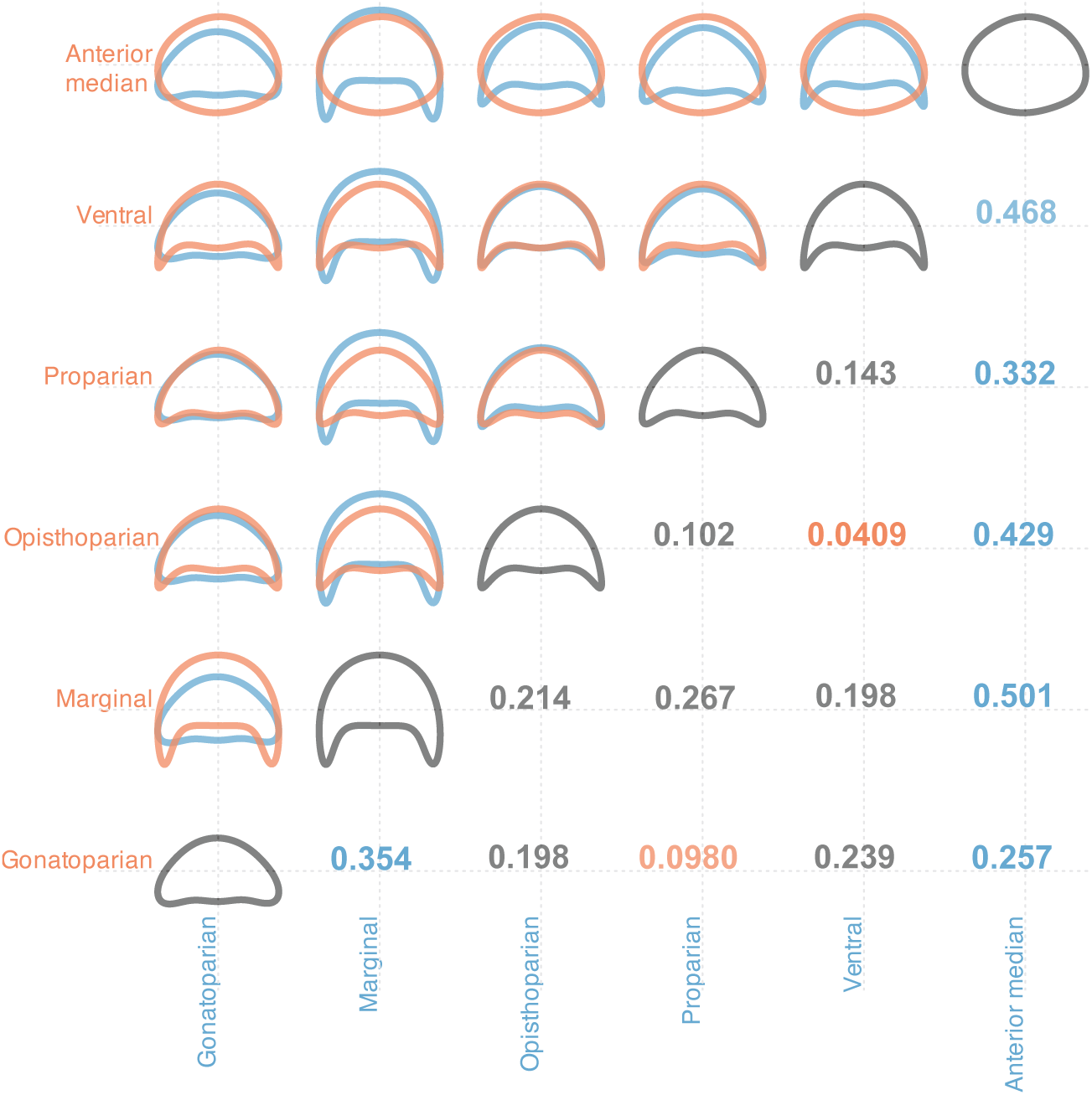
Pairwise comparison of average cephalic shape for the facial suture type groups, and pairwise centroid distances in Principal Components Analysis morphospace between the groups. Notably close centroid pairs (<0.1) are in orange text, and notably distant centroid pairs (>0.25) are in blue text.

As for the moulting mode groups, the LDA cross-validation table (Fig. 8B) suggests that cephalic outline shape cannot predict facial suture type. New species added to the dataset with facial sutures of any of the studied types are almost certain to be predicted as having opisthoparian facial sutures (all groups with a probability of >0.95). For all types (except opisthoparian), there is therefore an almost zero chance of a new species having its facial suture type correctly predicted.

The SoVs are notably low for the gonatoparian and anterior median facial suture groups (0.021 and 0.012 respectively; Fig. 10A), with pairwise t-tests (p = <<1×10^−20^) confirming this is significantly so (Supplementary III). This suggests low disparity, with the specimens tightly packed within morphospace for these two groups. For the gonatoparian group this evidently not a sample size issue (Table 1) but confirms the presence of outliers artificially extending the convex hull area and the SoR (1.233) of this group. For the anterior median group, the SoR is also low (0.407; Fig. 10B), though this result is likely to have been impacted by low sampling. The opisthoparian group has a moderate SoV and a high SoR (0.063 and 2.225 respectively; Fig. 10), indicating both species positioned towards the extremes of morphospace occupation and some clustering within this space, equating to a moderate overall disparity. The marginal suture group has both a high SoV and SoR (0.110 and 1.939 respectively; Fig. 10), suggesting disparate occupation of a large area of morphospace, and thereby high overall disparity. The proparian and ventral suture groups both show moderate SoV and SoR values. Almost every pairwise t-test for both the SoV and SoR results for facial suture type groups are significant (Supplementary III), suggesting true differences in the groups’ disparities.

**Figure 10:**
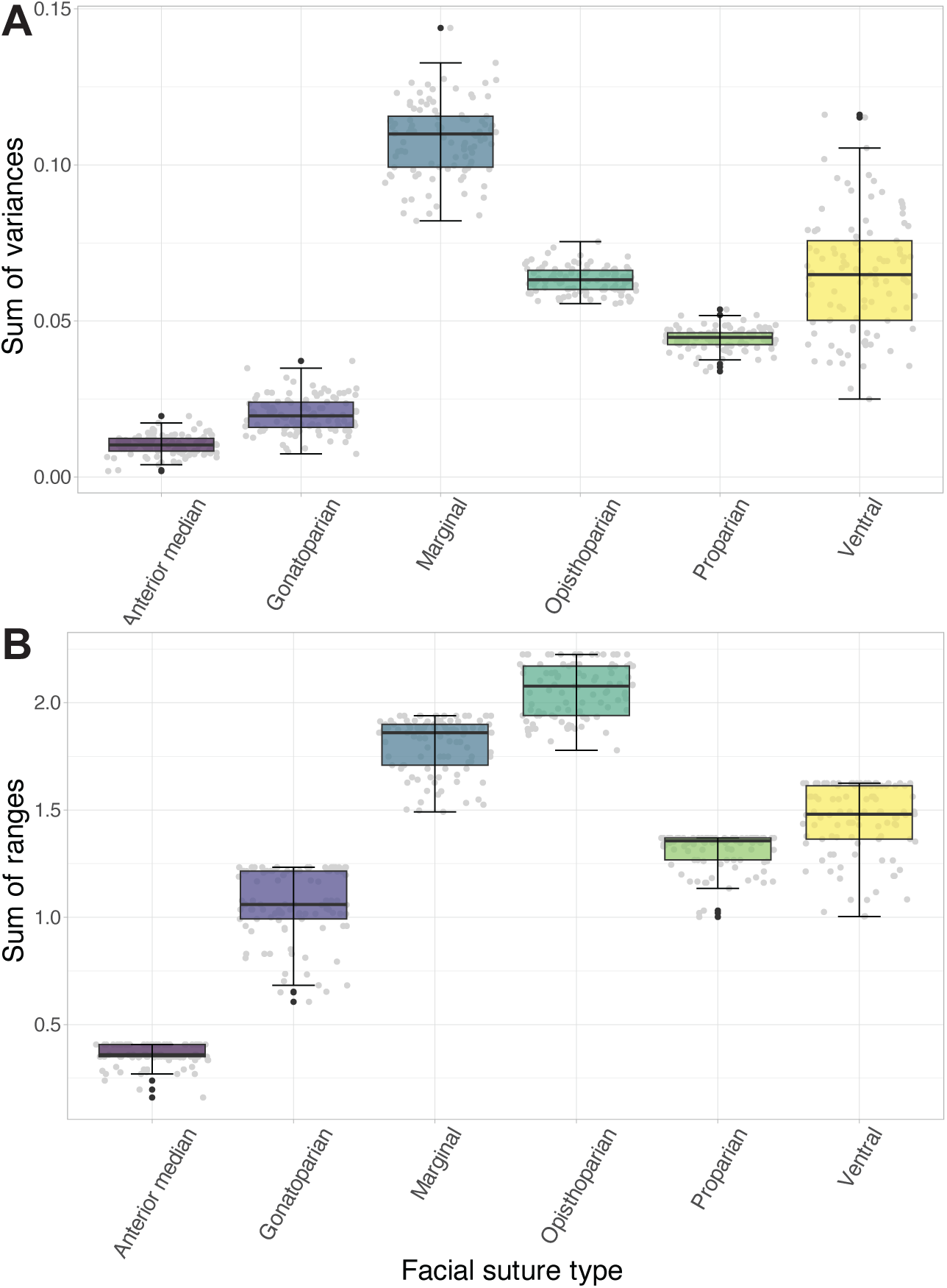
A, sum of variances box and whisker plots for facial suture type groups of the Principal Components Analysis; B, sum of ranges box and whisker plots of the same. Black lines show the median values, hinges correspond to the first and third quartiles, and the error bars represent 1.5× the interquartile range. Black dots are those species that fall outside the error bars (i.e., potential outliers), and the grey dots display all underlying data points (horizontal placement of points is meaningless).

Fitting of a linear regression model testing the linear response of PC position suggested only minor dependence between moulting mode and facial suture type. The estimates of change in PC1 for each moulting mode are generally small (almost all under 0.2; Supplementary III, Supplementary Figure 3), though three Salter’s moulting mode pairs are significant. This indicates that the PC1 positions of the Salter’s mode species are influenced by the type of facial suture the species have, but not the other moulting modes analysed (see Supplementary III). For PC2, all change estimates are very small (usually <0.1) and none of the t-tests were significant (Supplementary III), suggesting little impact of facial suture type on the PC2 position for the different moulting modes. Overall, the linear regression model demonstrates that facial suture type has a very minor impact on the cephalon outline shape of species showing the different moulting behaviours.

### Linear Mixed-Effects Models

Linear Mixed-Effects Models (LMMs) were created to determine the relative importance (effect sizes) of the categorical variables for determining cephalon outline morphometry (represented by PCA morphospace position). To test the impact of order assignment (as the fixed effect) on PC1 position, the best model (AIC = −465.2) is:

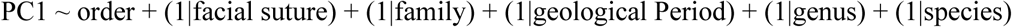

Thus, the model testing the effect of order assignment on PC1 position had its fit maximised by removing moulting mode entirely. The conditional R^2^ (cR^2^) states that 66% of the total variance in PC1 is attributable to this model, while the marginal R^2^ (mR^2^) states that 12% of the total variance in PC1 is attributable to the fixed effect of order assignment. Therefore, when adding a new species, we would be able to explain 12% of its PC1 position by order assignment. Order assignment has a significant impact on PC1 position (p = 0.008), with the effect size having a range of 0.13–0.02 (i.e., changing the order causes a maximum of 0.13 change in PC1 and a minimum of 0.02 in PC1). For the impact on PC2, the best-fitting model was the same as for PC1 but also had the facial suture type random effect variable removed (AIC = −1286.8 compared to −1284.8). Order assignment also has a significant impact on PC2 (p = <0.001), with an effect size of 0.47–0.03. This model explains 62% of total PC2 variance (cR^2^), with a notably high 32% of this variance being explained by order assignment (mR^2^). Belonging to the Harpida or Trinucleida is most influential on morphospace position based on the effect sizes for PC1 (Harpida = −0.52 to −0.01; Trinucleida = −0.44 to −0.10; Fig. 11E) and Harpida or Lichida for PC2 (−0.45 to −0.25 and +0.08 to +0.28 respectively; Fig. 11E); these effects are all significant, as are several other smaller order effect sizes (see Supplementary III). Belonging to the Asaphida, Phacopida and Redlichiida has close to no impact on PC2 (Supplementary Figure 4), aligning with the orders placements close to the centre of morphospace (Fig. 2), and for PC1 Aulacopleurida, Corynexochida, and Proetida assignment has almost no effect.

**Figure 11:**
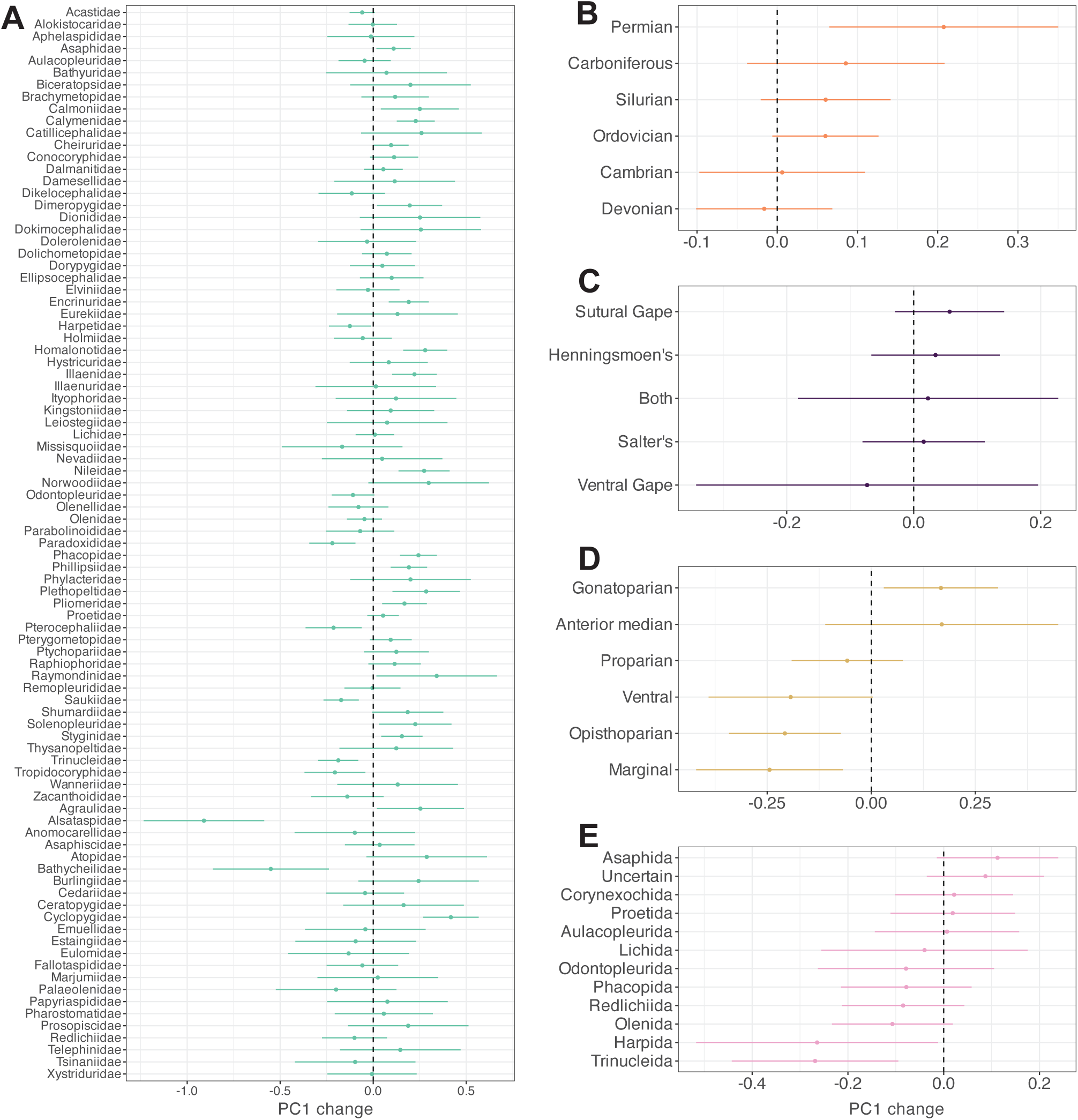
Best-fit Linear Mixed Models predictions for PC1. LMM fixed effects are: A, family; B, geological Period; C, moulting mode; D, facial suture type; E, order. Points are the average predictions of the fixed effect change of PC1 and whiskers represent the 95% confidence intervals. The LMM predictions for PC2 are available in Supplementary Figure 4.

To test the impact of geological Period assignment on PC1 position, the best model (AIC = −470.5) is:

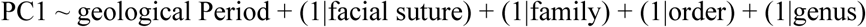

As for order assignment, the inclusion of moulting mode needlessly complicated the model, decreasing its fit, though model fit was also improved by removing species as a random effect. This model again explains 66% of the dataset PC1 variance, though the mR^2^ shows that only 4% of PC1 position variation is explainable by the fixed effect of geological period occupied. Despite being minor, this effect on PC1 is significant (p = 0.006) with a small effect size of 0.05− −0.02. Belonging to the Permian Period has the biggest impact on PC1 position, causing a 0.07−0.35 change, with this also being the only significant effect size (Fig. 11B; Supplementary III). For PC2 position, the best fitting model (AIC = −1265.8) is one with few random effects:

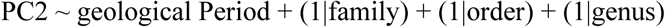

While this model explains 71% of the overall variance in PC2, the notably low mR^2^ at 4% shows that, as for PC1, very little of this variance is attributable to geological Period. Further, the fixed effect is only 0.04– −0.02, and this effect is non-significant (p = 0.065). This indicates that occupation of geological Period has no real impact on PC2 position of trilobite cephala.

To test the impact of family assignment on PC1 position, the best model is a very simple one, limited entirely to taxonomic variables:

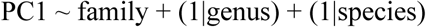

Like for the previous fixed effects, model fit for family as the fixed effect was improved by removing moulting mode, but also facial suture type. Unlike for the other models, removing geological Period very slightly improved the AIC value (−483.6 to −485.6), as did taxonomic order (same improvement, with including both variables increasing the AIC to −481.6), though the AIC was lower when keeping both species and genus as random effects. This is true also for the model estimating PC2 (AIC = −1293.5 to −1295.5 for both Period and order, −1291.5 if including both variables). The family fixed effect explains almost half of the variance in PC1 position at mR^2^ = 47%, while the model above with only four variables total itself represents 62% of the total dataset variance. The effect size range for family assignment on PC1 is large and significant at 0.88–0.01 (p = <0.001); the magnitude of impact family assignment has on PC1 thereby varies with the specific family. For example, being part of the Alsataspididae, Bathycheilidae and Cyclopygidae has a particularly strong impact on the value of PC1, with most other family assignments causing a less extreme change in PC1 of between around −0.25 and 0.25 (Fig. 11A; Supplementary III). The best-fitting model for PC2 is the same as for PC1 (AIC = −1295.5), and has a significant, large fixed-effect size of 1.05–0.02 (p = <0.001). Further, the model explains 64% of PC2 variance, and 51% of this variance is explained by family assignment. For PC2 position, belonging to the Harpetidae, Ityophoridae, Bathycheilidae and Telephinidae has the biggest impact (Supplementary Figure 4; Supplementary III).

To test the impact of facial suture type on PC1 position, the best model (AIC = −472.3) is:

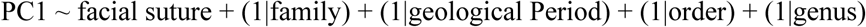

Facial suture type explains a small amount of the variance in PC1 position (14% mR^2^, 64% cR^2^), much more than geological Period and very slightly more than order assignment. The effect size of facial suture type on PC1 is significant (p = 0.001) at 0.17–0.05, with the marginal, opisthoparian, and gonatoparian suture types having significant effect sizes (Fig. 11D; Supplementary III). The PC2 best-fit model (AIC = 1264.5) also has geological Period removed:

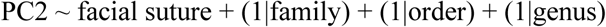

This model similarly explains 61% of the total variance in PC2, with 8% being due to facial suture type. The effect size is comparable to that for PC1 at 0.09– –0.25, but with this not being significant (p = 0.110).

To test the impact of moulting mode on PC1 position (in the subsetted dataset; see Methods), the best model incorporates a first-degree polynomial (linear equation; AIC = −105.5):

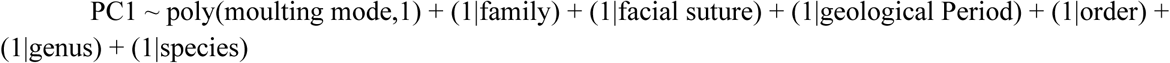

This model explains 87% of the overall variation in PC1, but moulting mode as the fixed effect explains none of this (mR^2^ = 0.00, effect size = 0.00, though p = 0.290). The best-fitting model without a polynomial term has a very slightly worse fit (AIC = −105.2; the same AIC is obtained whether the species variable was included or excluded) and excludes the geological Period variable. This model has the same cR^2^ value as for the model with polynomial, though with moulting mode explaining a tiny (and non-significant, p = 0.577) 1% of the PC1 variance (Fig. 11C). The higher cR^2^ values for the models with moulting mode as the fixed effect is due to the use of a subset of the larger dataset, including only species for which we had moulting data.

The best fitting model for PC2 is (AIC = −373.9):

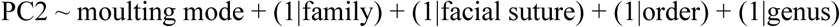

Unlike for PC1 position, moulting mode does explain some of the variation in PC2 position with a mR^2^ of 4% (and cR^2^ of 62%). The effect size of moulting mode on PC2 is weakly significant (p = 0.033) at 0.04–−0.04; moulting mode thereby had a minor impact on PC2 position (Supplementary Figure 4), but not PC1 position.

## 5 Discussion

### Can cephalon morphometry predict moulting behaviour?

The dataset demonstrates very little association between cephalon outline morphometry and exoskeleton moulting behaviour. There have been a number of studies quantifying the high levels of intra- and interspecific variability described for trilobites across the Palaeozoic (Brandt, 2002; Daley and Drage, 2016; Drage et al., 2018a, 2023; Drage, 2019, 2024; Corrales-García et al., 2020; Piccoli et al., 2021; Wang et al., 2021; Zong and Gong, 2022), and presenting various potential explanations for this variability. This includes suggestions that moulting variability might be related to evolutionary longevity (Brandt, 2002), ontogeny (Drage et al., 2018b), taxonomic group (Drage, 2019), thoracic segment count (Drage, 2024), and body size (Drage et al., 2023). A key aim of the study presented here was to determine whether a more complex morphological dataset had any relation to moulting behaviour, in particular the shape of the cephalon, which is presumed to be intrinsically related to moulting because all sutures related to moulting behaviour are part of, or proximal to, the cephalon (i.e., facial, rostral, hypostomal, ventral, marginal sutures, and cephalothoracic joint). However, the morphometric results presented here suggest very little association between moulting behaviour and cephalon outline morphometry, with almost complete nesting in morphospace (Fig. 4) and a low F value supporting little difference between the group morphospace positions (Table 1). The LDA results demonstrate almost no predictive ability of cephalon shape for moulting behaviour (Fig. 5B), further supporting a negligible association between the two. Lastly, the LMMs demonstrate moulting mode has almost no impact on position in morphospace, with moulting mode having no effect on PC1 position and a very minor effect size for PC2 (Fig. 11C), and most LMM models having improved fits by excluding the moulting variable.

Interestingly, the cephalon outline shapes of the facial suture and moulting groups were only minorly related based on the linear regression model (Supplementary III; Supplementary Figure 3), which suggests trilobite species did not moult significantly differently whether they had proparian, gonatoparian, or opisthoparian sutures. Most trilobite species moulted through a sutural gape opened at the facial sutures (Henningsmoen, 1975; Drage, 2019). Several suture groups (marginal and anterior median) showed very divergent average cephalon shapes (Fig. 9), though from a moulting perspective species with these suture types still used the Sutural Gape or Ventral Gape modes of moulting. Overall, there seems to be little utility in considering facial suture (particularly dorsal) type for explaining moulting behaviour, except for the presence or absence of facial sutures leading to an ability (= Sutural Gape mode) or inability (= Salter’s mode) to use them to create a moulting sutural gape. More can only be said about moulting behaviour when the suture type of the species is known, and only then if the suture type is specific to a certain moulting behaviour (not the different types of dorsal suture); that is, taxa with ventral sutures (e.g., Olenellidae) moult ventrally, and those with marginal sutures (e.g., Harpididae) moult using a marginal suture gape.

Overall, the results here suggest variability in trilobite moulting behaviour cannot be explained by cephalon shape. Coupled with the findings of previous studies that found no clear statistical associations between moulting variability and traditional morphometry (Drage et al., 2023; Drage, 2024), this work supports that trilobite moulting behaviour is relatively phenotypically plastic. This means that, for most of Trilobita, variables potentially related to moulting behaviour cannot be identified on a general basis, excepting for groups with highly specialised moulting structure adaptations, like Harpetida. It is probable that, as for most animal behaviours, there are too many underlying variables complicating the end phenotype, and that any explanation is relevant only at the small-scale, such as species-level associations with ontogenetic trajectories and morphological changes (Drage et al., 2018b). Additionally, quantitative studies of arthropod moulting patterns in deep time have yet to analyse the impact of preservational biases, like biostratinomic processes, on the preservation of moult assemblages and the resulting interpretations of plasticity. This does not mean, however, that we should not continue to study trilobite moulting within species, genera, or families, as moulting remains a crucial factor in the life cycles of trilobites, and is intrinsically linked to their growth and development.

### Cephalon outline shape is best explained by taxonomic assignment

Of all the variables investigated, taxonomic assignment best explains the variation in cephalon outline morphometry in the dataset. This is clearest in the LMM results, with order assignment explaining 12% of variability in PC1 (32% in PC2) and family 47% of variability in PC1 (51% in PC2). The LMM with family as the fixed effect had the best fit when including only taxonomic variables as the random effects, suggesting that these variables are far more impactful as predictors of morphospace position than the non-taxonomic variables (geological Period, facial suture type, moulting mode). This is a reassuring finding given trilobite taxonomy is highly based on cephalon morphology (both due to cephalon morphological disparity and greater distinctiveness than thoracopyga) and the cephalon hosts many key behavioural and sensory structures (e.g., Clarkson et al., 2006; Edgecombe and Fortey, 2023), so we would expect a strong correlation. However, the better fit of this model when excluding the order variable indicates that order assignment might actually be interfering with the ability of family assignment to model cephalon shape; that is, there is a lack of consistency between the order and family cephalon outline predictions. This finding provides quantitative evidence that higher level trilobite taxonomy remains less reliable and family or lower-level taxonomy better captures trilobite morphological variation and evolutionary relationships (Adrain, 2011; Paterson, 2019).

Additionally, it is interesting that many best-fit LMMs had species removed as a random effect (though not genus), including those with geological Period, facial suture type, and moulting mode as the fixed effect, but not order and family assignment. However, genus was strongly influential as a random effect, with its removal greatly decreasing the AIC fit of models. This suggests that trilobite genera strongly differ in their cephalic shape, but the shape difference within these genera (i.e., between species) is generally not significant. It is possible that this result is affected by more than half of the genera in the dataset being monospecific (250 monospecific genera out of 402 total), so that there is little additional variation explainable by species that is not already explained by genus.

The family results are also interesting on a finer scale, with differences in clustering patterns across PCA morphospace that might be informed by ecological aspects. For example, the Cyclopygidae are extremely clustered at the far positive end of PC1 (Fig. 2) as they have a uniquely rounded average cephalon outline (Supplementary Figure 1), which may be explained by their suggested (meso-) pelagic ecology based on morphology but also primary occurrence in deep-water facies (Fortey, 1981, 1985, 2014). However, species of Remopleurididae have also been suggested to be pelagic (Dollo, 1910; Marek, 1962), largely as a result of having similar overall morphologies to cyclopygids (Bergström, 1973), but they show less clustering in morphospace and a more typical average cephalon shape (Fig. 2; Supplementary Figure 1). This finding agrees with the conclusion of Fortey (1985), who notes it unlikely that remopleurids were pelagic based on various morphological aspects, lack of streamlining, and general absence in deep-water facies. Telephinidae were also putatively (epi-) pelagic (Fortey, 1981, 1985), independently arising to in the groups already discussed, and here show a unique mean cephalon shape owing to large eyes and spines (Supplementary Figure 1), though we have not presented their placement in morphospace due to low sampling in the dataset.

A few groups display multiple distinguishable clusters within the breadth of their morphospace occupation. In particular, the Dalmanitidae shows two clusters with a comparable spread of PC2 values but one at positive PC1 values and one at negative (Fig. 2). The Acastidae also show a somewhat similar pattern of two clusters either side of the centre on PC1. For the Dalmanitidae, these clusters may correspond with the group’s different subfamilies (Holloway, 1981); this might be possible to visualise if plotting the subfamilies separately in morphospace, however, the subfamilial assignments of Dalmanitidae taxa are often not resolvable because the species were described many years ago, and the relationships of the family have not been updated and need treatment (Rustán and Vaccari, 2012). The positive PC1 dalmanitid cluster occupies a comparable position to the strongly clustered Phacopidae, another highly diverse family within the order Phacopida. The phacopid families show quite dense clustering close to the centre of morphospace and usually slightly positive on PC1 (Fig. 2). This suggests that the order did comparably little exploration of morphospace in any of its families (excepting the Cheiruridae, occupying a notably higher area of morphospace), potentially also indicating comparable morphologies and ecologies across the group (i.e., their large holochroal or schizochroal eyes, this similar cephalic shape; Supplementary Figure 1), potentially with avoidance of competition through geographical and temporal spread. With both having extremely long histories of research focus, high taxonomic diversity, and large sample sizes it is interesting that the Proetidae occupy a larger area of morphospace and have a higher disparity than the Phacopidae (Figs 2, 3). The two encompassing orders (Proetida and Phacopida) do seem to show contrasting patterns of diversity, disparity, and environment across the Palaeozoic (Hopkins, 2014; Drage, 2019; Bault et al., 2022; Drage and Pates, 2024).

Other families show little obvious clustering, containing species that are more evenly spread across the extents of their convex hulls (excepting potential outliers; see Results), and often overlapping the convex hulls of other families. This indicates that, while families might be restricted to a subset of the total area trilobites occupied, the species within most families show disparity in cephalic shape. For example, the families comprising order Asaphida have convex hulls more spread across the total area of occupied trilobite morphospace, but were also taxonomically diverse like phacopid families. Asaphida encompasses a diversity of families living across major periods of global diversification in the Cambrian and Ordovician (Drage and Pates, 2024), and, while Asaphida are often considered ‘conservative’ trilobites, this includes some families with divergent ecologies. For example, the putatively pelagic Cyclopygidae (Fortey, 2014) and probable infaunal groups like Nileidae (Fortey, 1986; Esteve et al., 2021b); perhaps explaining this greater morphometric disparity than other highly diverse groups like Phacopida. Despite being trilobites with unusual morphologies, being notably spiny often with exoskeleton tubercles, the Lichidae occupy a reasonably large area of morphospace, with many species occurring within the main centre of morphospace occupation. Interestingly, Hopkins et al. (2023) did find that the Lichida was the only major trilobite order living in the Devonian still evolving novel morphological character combinations, which agrees with their high morphospace area and occupation of the extreme of space at the most positive of PC2 (Fig. 2).

The family Olenidae occupy the largest area of morphospace of all families, while also showing little clustering of species within their convex hull (Fig. 2). Some Olenidae have been suggested to be chemoautotrophic symbionts, using a symbiotic relationship with sulphur-consuming bacteria to feed (Fortey, 2000, 2014). This is owing to their frequent preservation in deep-water, probably low oxygen, shale facies (often termed ‘olenid biofacies’), dorsoventrally flattened bodies with many thoracic segments, thin exoskeleton, and reduction of hypostome in some taxa (Clarkson and Taylor, 1995; Fortey, 2000, 2014). However, this hypothesis of conserved morphological adaptations for a specialised habitat suggests that olenid cephala would be fairly low in disparity (Fortey, 2000), which is not the case based on this dataset. This finding supports contrasting work that found olenids to have had a broader spread across environments differing in bathymetry and oxygen availability, rather than being restricted to this traditional chemoautotrophic, deep-water view (Balseiro et al., 2011). Indeed, even Fortey (2000), who argued that the Olenidae as a whole were chemoautotrophs, noted that there is morphological variability within the Olenidae that suggests adaptations to other ecologies, and the disparity results here support this suggestion. Lastly, like many higher level trilobite taxonomic groups, Olenidae has been suggested to not be monophyletic, with analyses by Monti and Confalonieri (2019) indicating several groups should be removed from the Olenidae, and this would be expected to reduce the group’s disparity.

Several families show notable contrasting patterns in their occupation of morphospace (Fig. 2). Harpetids and trinucleids are frequently considered together as ‘trinucleimorphs’ during studies of trilobite palaeoecology, with comparisons drawn between their broad and deep cephala with long genal spines, frequently horseshoe-shaped marginal brims, and cephalic pitting (Fortey and Owens, 1990; Fortey and Gutiérrez-Marco, 2022; Beech et al., 2024; Pates and Drage, 2024). This cephalon morphology is thought to be convergent within the two groups (and others), representing adaptations to comparable ecologies, though their actual function is still debated (e.g., suspension feeding, preventing sinking into the sediment, strengthening, etc.; Fortey and Owens, 1999; Pearson, 2017; Fortey and Gutiérrez-Marco, 2022; Beech et al., 2024; Pates and Drage, 2024). The relevant orders (Harpida and Trinucleida) had the strongest effects on PC1 and PC2 placement in this dataset, according to the best-fitting LMM (Fig. 11E), so have notably divergent cephalic shapes from the rest of Trilobita. Despite this, the two groups’ cephalon outlines are more internally disparate than might be expected for apparently being adapted to a specialised, unusual life mode, with the groups having reasonably high SoVs (low clustering within area occupied), SoRs and morphospace occupation areas (Figs 2, 3). Despite their supposedly comparable cephalic adaptations, the two families occupy very different areas of morphospace with almost no overlapping (Fig. 2); Harpetida are slightly negative on PC1 but at the positive extreme of PC2 (except for one, likely anomalous, specimen), and Trinucleida are spread across a reasonably wide area close to the negatives of both PC1 and 2. The similarity between harpetid and trinucleid cephala may therefore be superficial, as is revealed in the average cephala; the harpetid cephalon is axially broader with broader genal spines (Supplementary Figure 1). Raphiophoriidae, belonging to the Trinucleida along with Trinucleidae (Bignon et al., 2020), have also been linked to trinucleimorphs, having been suggested to have fed comparably (Fortey and Owens, 1999), or used their long cephalic spines for comparable sensory or sexual selection purposes (Knell and Fortey, 2005; Vannier et al., 2019; Gishlick and Fortey, 2023). However, raphiophoriids occupy a large and high-disparity area of morphospace that only slightly overlaps with the Trinucleidae at the centre of morphospace (Fig. 2). Some raphiophoriid species do have comparable cephalon shapes with long spines, but their average cephalon is more similar to that of other trilobite families (rather than the divergent harpetid and trinucleid cephala). This is presumably owing to their morphological variability (Fortey, 1975) rather than single-shape specialism and potentially related to the paraphyly suggested by Bignon et al. (2020). These results suggest a range of cephalon shapes could have been adaptations to a comparable trinucleimorph ecology, whether this represents adaptation to suspension feeding, the snowshoe effect, or another function. Conversely, this may also indicate that these groups, while having superficial morphological similarities, are adapted to different ecologies and that the cephalic brim and long spines did not fulfil comparable functions.

Many families had an extensive effect of PC1 position, as predicted by their effect sizes in the best-fitting LMM not overlapping PC1 = 0 (Fig. 11A). This particularly includes the Alsataspididae (−ve PC1), Bathycheilidae (−ve PC1) and Cyclopygidae (+ve PC1). For the Alsataspididae, the mean cephalon shape is clearly extreme, with long, narrow genal spines and an anteriorly directed cephalic spine (Supplementary Figure 1). The Bathycheilidae also have particularly long genal spines, and the Cyclopygidae a rounded cephalon that is likely related to a potential pelagic ecology, as discussed above. The Alsataspididae and Bathycheilidae are less well-sampled in this dataset, with the former having problematic taxonomy within Trinucleida (Bignon et al., 2020), though with both families likely having this strong impact on morphospace placement for the same reason as the Trinucleidae (both belonging to the Trinucleida with long spines suggested to prevent sinking into sediment; Hammann, 1985). However, 29 families in total had confidence intervals that did not span PC1 = 0 (Fig. 11A). These family assignments had particularly strong effects on the placement of species in morphospace, suggesting these taxonomic groups are well-defined based on the cephalon outline shape.

The stratigraphically earliest families, in Cambrian Stage 3, are the Fallotaspididae, Bigotinidae, Abadiellidae, and Holmiidae (Zhang et al., 2017). This dataset only includes Fallotaspididae and Holmiidae, with these families having little effect on PC1 placement in the LMM (Fig. 11A), and holmiids largely occurring at the centre of occupied morphospace excepting two species (Fig. 2). Both families belong to the order Redlichiida and suborder Olenellina. The other early Cambrian family Olenellidae is part of the same suborder, and the similarly lower Cambrian Ellipsocephalidae and Paradoxididae the same order. Most of these species also show central positions in morphospace (Fig. 2), relatively low disparity measures (Fig. 3), and, except for the Paradoxididae, have low LMM effect sizes as well as a relatively low order (Redlichida) effect size (Fig. 11). This supports the supposition that trilobites originated in a more constrained area of morphospace under less variety of environmental and ecological pressures (e.g., Hopkins, 2014) and increasing in disparity more so after the early Cambrian than during their initial diversification, reflecting the Cambrian evolutionary rate stasis noted by Paterson et al. (2019). Smith and Lieberman (1999) even found no real change in constraint or flexibility in olenelloid trilobites over this earliest period of their evolution. However, this stasis may be limited to the cephalic outline, with variation occurring in parts of the morphology not captured by our analyses. For example, Holmes (2023) found a rapid increase in cephalic disparity when focusing on a much more granular timeframe in the early Cambrian and incorporating a greater variety of cephalic morphometric data, and Webster (2007) found the highest levels of phenotypic polymorphism in the early Cambrian, again considering a greater range of morphology than just cephalic outline. In contrast, families living later in the Palaeozoic (e.g., Proetidae, Phillipsiidae, Aulacopleuridae), particularly during the Carboniferous and Permian, do not necessarily show more restricted occupation of morphospace as expected by their decline in diversity towards their eventual extinction. However, their position is towards the centre of morphospace in areas occupied since the early Palaeozoic (Fig. 2), as also found by Hopkins et al. (2023), though we did not find (unlike Hopkins et al., 2023) that the Proetidae showed notably low disparity compared to other families.

### Other predictors of cephalon outline shape

Geological Period explained very little of the dataset variance in PC1 and 2 (Fig. 11B, Supplementary Figure 4; see Drage and Pates, 2024, for PCA morphospaces). However, assignment to the Permian did have a positive, significant effect size for PC1 (Fig. 11B), unlike all other Periods. This presumably results from the consistent restriction to the centre of morphospace (and slightly negative on PC2) of the Permian groups (Proetidae and Phillipsiidae, order Proetida; Fig. 2), though as discussed above their disparity is not necessarily lower than many earlier families. However, there remains finer-scale trends to consider in the lead up to the end-Permian extinction of Proetida and trilobites as a whole, such as their increased range restriction and endemism (Brezinski, 2023).

Facial suture type had the third-greatest impact on PC1 and 2 variances after family and order, as modelled by the LMMs (Fig. 11, Supplementary Figure 4). Cephalic shape alone was incapable of predicting facial suture type (Fig. 8B), though facial suture had a reasonably high effect size and the suture types gonatoparian (+ve PC1), ventral (−ve PC1), opisthoparian (−ve PC1) and marginal (−ve PC1) all had effect sizes with confidence intervals not overlapping PC1 = 0 (Fig. 11D). For most of these, the facial suture types are somewhat restricted to specific trilobite taxonomic groups (being amongst their diagnosable characters), likely explaining this result. For example, gonatoparian sutures are largely found in the phacopid suborder Calymenina (e.g., the families Calymenidae, Homalonotidae), marginal sutures usually in the Harpida, and ventral sutures usually in the stratigraphically earliest redlichiid trilobites. However, the result for the opisthoparian suture grouping is interesting, as this is the most widely spread general suture type in trilobites, being found across most orders from the early Cambrian (Fortey, 2001).

Other ecological factors, such as feeding mode and life mode, may also impact cephalic morphometry. For the most part, hypotheses of ecological mode remain quantitatively untested, albeit frequently based on multiple lines of morphological and contextual evidence (Fortey and Owens, 1999; Fortey, 2014). Some trilobite species have seen more detailed ecological hypothesis-testing (e.g., Pearson, 2017; Esteve et al., 2021a; Esteve and López-Pachón, 2023), though many suggestions of ecological mode remain to be tested. This means that general trilobite ecological data (e.g., from the PBDB; Uhen et al., 2023) is not sufficiently detailed to test broad-scale life and feeding mode patterns, either being imputed from high taxonomic levels (meaning it is not relevant to actual species’ morphology) or not having meaningful differences. These available ecological data (i.e., hosted on the PBDB) might be of sufficient detail for other extinct animal groups, but trilobite ecological hypothesis-testing needs more attention before these variables are usable in broad-scale analyses (see Supplementary IV).

Finally, the results presented here are partially a reflection of the methods used. Morphological information is lost when comparing groups using geometric morphometrics; this necessitates comparing structures across clades that we are confident are biologically homologous or otherwise directly comparable (Collins et al., 2021). This means we must make compromises when using geometric morphometrics, particularly when including taxa across broad taxonomic groups, as here, such as analysing only semilandmark outlines, or landmarking of the few homologous structures of which we can be confident. It is likely that this reduction in usable morphological information contributes to the lack of predictive ability we see in these morphometric analyses, though this is presumably less of an issue in Trilobita than in groups with high three-dimensionality and uncertainty in homology, such as for Gastropoda (see Collins et al., 2021).

## 6 Conclusions

We presented associations between trilobite cephalic outline morphometry and family-level taxonomy, facial suture type, and moulting behaviour, from a dataset of 762 species (Suárez and Esteve, 2021; Drage and Pates, 2024) across the Palaeozoic, as well as the first exploration of the hierarchical importance of different factors affecting trilobite morphometry using Linear Mixed-Effects Modelling. Following the results of other analyses exploring the drivers of trilobite moulting behaviour variability (Drage et al., 2023; Drage, 2024), we found that moulting variation cannot be linked with any certainty to cephalic shape, nor to facial suture type. Based on this dataset, moulting behaviour cannot be predicted from cephalic shape. These results, coupled with previous quantitative studies of trilobite moulting, suggest that variability on this broad scale is largely unexplainable with any real certainty probably due to a combination of overall presence/absence of facial sutures (and specialised sutures like ventral or marginal sutures), the randomness of whether the cephalothoracic joint opens as well as cephalic sutures when pressure is applied to the exoskeleton during moulting, and the influence of post-moulting abiotic processes potentially causing additional disarticulation or movement of sclerites. Moulting remains a crucial factor in understanding the life histories and evolutionary trajectories of extinct and extant arthropods, so, despite these results, should remain a priority for study.

The results of this study also demonstrate the utility of Linear Mixed-Effects Models for palaeontological research, to determine the hierarchical importance of different variables on fossil data. These LMM analyses elucidated the drivers of trilobite cephalic shape, supporting that family-level assignment is, overall, stable and well-explains cephalic morphology, and more so than order assignment, in which the orders significantly overlap in cephalic shape (Drage and Pates, 2024). These LMMs also supported the lack of influence of certain variables on cephalic shape, notably moulting behaviour (as discussed above), geological Period of occupation, and facial suture type. This study highlights the use of LMMs for interrogating broad-scale fossil morphological data to unpick the effects of different variables in what are inherently complex, multi-variable systems.

## 7 Acknowledgements

We are grateful to Jana Bruthansová (National Museum Prague, Prague), Richard Howard and Zoe Hughes (Natural History Museum UK, London), Matt Riley (Sedgwick Museum, Cambridge), Mónica M. Solórzano Kraemer (Senckenberg Museum, Frankfurt), and Julien Kimmig (Staatliches Museum für Naturkunde Karlsruhe) for providing access to specimens at their respective institutions. In addition, we thank all the museums, volunteers and staff who uploaded images of trilobites to iDigBio, which facilitated data collection for this research during periods of remote working during early 2021. HBD was funded under a Swiss National Science Foundation Sinergia grant (198691). SP was supported for this work by a NERC IRF NE/X017745/1 (University of Exeter).

## 8 Author contributions

**Harriet B. Drage:** Conceptualisation, Methodology, Software, Validation, Formal Analysis, Investigation, Data Curation, Writing – Original Draft, Writing – Review & Editing, Visualisation, Project Administration. **Stephen Pates:** Conceptualisation, Methodology, Software, Validation, Formal Analysis, Investigation, Data Curation, Writing – Review & Editing, Project Administration.

## 10 Supplementary files

All supplementary files are available open access on the Open Science Framework at https://doi.org/10.17605/OSF.IO/EH86Z

**Supplementary folder I:** Data files for all analyses. Folder contains the raw data used, with entries reduced to a single incidence of each species; HDSPdata_raw_DP25.csv is the original data collected by the authors and first published in Drage and Pates (2024), SEdata_raw_DP25.csv is the data used here and first published in Suárez and Esteve (2021). totaldata_PCs_DP25.csv is the processed principal components coordinates used for all analyses (PCs 1 to 10); this is the two raw data files combined and with the Suárez and Esteve (2021) data resampled to give equivalent numbers of data points, combined with the final data for all other variables, subject to elliptical Fourier transformation, and converted to a data frame; all steps to produce this file are given in the R code in Supplementary II. totaldata_counts_DP25.csv gives the counts for each group of each variable in the total dataset.

**Supplementary II:** R code used for all data processing and analyses; Supp2_DP2025_code.R.

**Supplementary folder III:** Supplementary result files. disparitytestdata_DP25.xlsx contains the results for all pairwise t-tests on the calculated disparity measures (sums of variances and ranges). linearregression_suturemoult_DP25.xlsx gives the full results of the linear regression analysis between facial suture type and moulting mode for PC1 and 2 (displayed in Supplementary Figure 3). LMMresults_DP25.xlsx give the full predictions of the best-fitting Linear Mixed-Effects Models for each variable taken as the fixed effect, for each of PC1 and 2. familycentroiddistances_DP25.csv gives the pairwise centroid distances for all family groups (regardless of sample size).

**Supplementary IV:** Description and PCA morphospace test results for interpreting cephalic outline morphometry using ecological and feeding mode. These tests were not included in the full study, as explained in this supplementary; Supp4_ecologytest_DP25.doc.

**Supplementary Figure 1:** Average cephalon shape for each family. All families are displayed, even those with n=<5, though these were not included in other downstream analyses. Colours correspond to the order each family belongs to.

**Supplementary Figure 2:** Linear Discriminant Analysis cross-validation table. The table gives the probability that a species with a given family assignment (y-axis) will be predicted as each of the families along the x-axis. Empty cells and missing pairings equate to a negligible (close to zero) probability. All families are displayed, even those with n=<5, though these were not included in other downstream analyses.

**Supplementary Figure 3:** A, Linear regression model estimates for changes in PC1 for species belonging to a moulting mode group and having different facial suture types; B, the same, but estimates for PC2 change.

**Supplementary Figure 4:** Best-fit Linear Mixed Models predictions for PC2. LMM fixed effects are: A, family; B, geological Period; C, moulting mode; D, facial suture type; E, order. Points are the average predictions of the fixed effect change of PC2 and whiskers represent the 95% confidence intervals.

